# Technical and biological variations in the purification of extrachromosomal circular DNA (eccDNA) and the finding of more eccDNA in the plasma of lung adenocarcinoma patients compared with healthy donors

**DOI:** 10.1101/2024.03.05.583481

**Authors:** Egija Zole, Lasse Bøllehuus Hansen, János Haskó, Daniela Gerovska, Marcos J. Araúzo-Bravo, Julie Boertmann Noer, Yonglun Luo, Jakob Sidenius Johansen, Birgitte Regenberg

## Abstract

Human plasma DNA originates from all tissues and organs, holding the potential as a versatile marker for diseases such as cancer, as fragments of cancer-specific alleles can be found circulating in the blood. While linear DNA has been studied intensely as a liquid biomarker, the role of circular circulating DNA in cancer is more unknown due, in part, to a lack of comprehensive testing methods. Our developed method profiles extrachromosomal circular DNA (eccDNA) in plasma, integrating Solid-Phase Reversible Immobilization (SPRI) bead purification, the removal of linear DNA and mitochondrial DNA, and DNA sequencing. As an initial assessment, we examined the method, biological variations, and technical variations using plasma samples from four patients with lung adenocarcinoma and four healthy and physically fit individuals. Despite the small sample group, we observed a significant eccDNA increase in cancer patients in two independent laboratories and that eccDNA covered up to 0.4 % of the genome/mL plasma. We found a subset of eccDNA from recurrent genes present in cancer samples but not in every control. In conclusion, our data reflect the large variation found in eccDNA sequence content and show that the variability observed among replicates in eccDNA stems from a biological source and can cause inconclusive findings for biomarkers. This suggests the need to explore other biological markers, such as epigenetic features on eccDNA.

## 1. Introduction

Until recently, the diagnosis and monitoring of many cancer types have almost exclusively relied on the usage of biopsies, contrast fluid and clinical signs. This has changed with an increasing scientific focus on developing safer and less invasive diagnostic approaches for cancer detection and monitoring (1). One such developing field focuses on liquid biopsies, among which plasma biopsies offer an easily applicable standard clinical approach, with low invasiveness for patient monitoring through novel biological markers (reviewed in (2)). Plasma contains cell-free DNA (cfDNA), which can serve as a biomarker for multiple diseases and conditions, including cancers (reviewed in (3)). Among cancers, lung cancer has the highest yearly death toll, and despite recent advances in early detection and treatment, the majority of cases are diagnosed at a late stage (4,5). Previous investigations of linear cfDNA in plasma from patients with cancers such as hepatocellular, colorectal, and lung cancer, have found copy number and size variations in linear cfDNA compared with healthy controls (6,7). Furthermore, it has been suggested that both the specific sequences, mutations and methylation profiles observed in cfDNA can serve as indicative biomarkers of cancer (8,9). In line with this, there are already FDA-approved linear cfDNA testing methods for specific alleles (reviewed in (10)), indicating the potential of cfDNA as a tool in disease diagnostics and monitoring.

However, the field of cfDNA is still developing, and cfDNA assays often show insufficient sensitivity and specificity for many cancers, especially for those in the early stages (8,11). Linear DNA is unstable (T1/2 = 114 min, (12)) and is only found at low concentrations in plasma, causing high variability of cfDNA concentration and content among patients and samples (13–19). This reduces linear plasma DNA’s applicability as a broad cancer marker despite its relevance as a direct indicator of tumor constituents. Another potentially interesting cancer biomarker is extrachromosomal circular DNA (eccDNA), which has the potential to contain genetic information similar to that of linear DNA. The circular form of eccDNA is likely to make it more resistant to enzymatic degradation and gives it increased stability in the bloodstream, compared with the linear DNA, as there are no free ends for DNA exonucleases to degrade (20,21). The potential of eccDNA as a diagnostic biomarker has not gone unnoticed by the scientific community, which has tested its potential in relation to the detection of cancer and fetal eccDNA (21,22). EccDNA is of particular interest as a cancer marker as it has been shown to be associated with both DNA damage (23–25) and tumor heterogeneity (reviewed in (26)). However, the methods used to extract cell-free eccDNA have not been standardized, and the current methods are considered to be time-consuming and lacking sufficient sensitivity (21,26,27). Methods for eccDNA extraction can also be DNA degrading, which can affect the purification yield of larger eccDNA (27,28). Since large eccDNA has the potential to contain valuable information, such as whole- or fragments of oncogenes (reviewed in (26)), purification that preserves the circular DNA is important for clinical assessments. EccDNA has previously been extracted from plasma samples of patients with lung cancers (21,29,30). EccDNA in these studies ranged in size from 100 bp to 400 bp (30), and 100 bp to 2000 bp (21,29). Kumar et al. show that two out of four lung cancer patients carry larger circulating circular DNA pre-surgery than post-surgery (21). Two groups have found an overrepresentation of eccDNA from particular genes in the plasma of adenocarcinoma patients, though there is no overlap between the genes in the two studies (29,30). While these studies suggest that eccDNA has potential as a marker for lung adenocarcinomas by investigating the variation in eccDNA between both patients and independent samples, they also reveal a need for methods that preserve the eccDNA.

We therefore developed a method for extracting eccDNA from plasma based on Solid-Phase Reversible Immobilization (SPRI) bead purification. It takes advantage of solid-phase reversible immobilization on magnetic beads and enzymatic reactions to remove mitochondrial DNA (mtDNA) and linear chromosomal DNA fragments from plasma samples of variable volumes in the mL range. EccDNA extracted by this method can subsequently be sequenced, mapped, and analyzed, including the eccDNA profiles for overrepresented genes. We tested our eccDNA purification method by comparing the results of eccDNA extracted from one mL plasma from four controls and four patients with stage IV lung cancer (adenocarcinoma). We performed purification at two different laboratories to see if results were reproducible in laboratories with different experience levels of the technique (Laboratory A was familiar with the method, whereas Laboratory B was not). To further understand the level of biological and technical variance, we measured the variation at different purification steps, from DNA extraction to the synthesis of sequencing libraries. Our study revealed that individuals with stage IV lung cancer could be distinguished from healthy individuals based on the number of eccDNA in their plasma. We also found that the number of circular DNA per sample varied between biological and technical replicates. In addition, the size of purified DNA was much larger than in previous studies. Although circular DNA profiles had little sequence overlap, a subset of circular DNA from genes involved in lung cancer was overrepresented in the individuals with lung cancer. Thus, the SPRI bead purification method appears to be suitable for the purification and application of plasma eccDNA. Still, for specific cancer biomarkers, future studies have to explore additional possible markers like epigenetic or ATAC-seq profiles on eccDNA.

## 2. Materials and Methods

### 2.1. Plasma samples

For the preliminary assessment of the method, we used two distinct groups of plasma samples of 2-6 mL per patient (1 mL per replicate sample) from 4 patients with lung cancer (adenocarcinoma, stage IV) (age 67.5 ±6.2 years, 1 male and 3 females). Samples were well mixed by gentle pipetting, avoiding high-speed centrifugation and vortexing to preserve circular DNA. Samples were obtained from LUCAS Biobank, and a group of 4 healthy and physically fit controls (age 57.5 ± 3.5 years, 1 male and 3 females), obtained from voluntary donors. The cancer patients’ blood was collected 11-27 days after the pathology diagnosis. The blood samples were collected before the start of chemotherapy treatment. The blood was collected in EDTA-containing BD Vacutainer tubes (Becton, Dickinson and Company, NJ, USA), from which plasma was separated by centrifugation at 2000 g for 15 min at room temperature and stored at - 80 °C until analysis. All plasma samples were anonymized. One of the cancer plasma samples from laboratory B (B6) failed during Φ29 polymerase rolling circle amplification, therefore, B6 was removed from further bioinformatics data analysis though still sequenced and applied as a control for non-Φ29 amplified eccDNA identification.

### 2.2. Plasmid and linear DNA controls

For quality control and procedure testing, we spiked-in a control mixture consisting of plasmids (50,000 copies of p4339 (5064 bp (base pairs)), 10,000 copies of pBR322 (4361 bp) (New England Biolabs, MA, USA)) (31), and amplified linear DNA fragments from yeast DNA (25,000 copies of linear DNA formed from primers detailed in **Table S1** that fit the four yeast genes GNP1, AGP1, ACT1 and BCP1). All plasmids were maintained in Escherichia coli and purified with a standard plasmid midi-prep kit (NucleoBond® Xtra Midi, MACHEREY-NAGEL, DE).

### 2.3. Circular DNA extraction by phenol/chloroform method

We employed a conventional phenol/chloroform-based salt precipitation method to extract DNA from six plasma samples, for the purpose of comparison with the SPRI bead purification method applied to another set of six samples. The tested plasma consisted of pooled human plasma from Innovative Research (MI, USA). In short, the phenol/chloroform purification approach was conducted as follows: for each replicate, 1 mL of plasma was used. For internal control, 10 μL of the spike-in mix was used. After spike-in addition, 22 μL of proteinase K (20 mg/mL) (ThermoFisher, MA, USA) was added to each sample (final concentration 400 μg/mL), followed by 64 μL 20% SDS (ThermoFisher, MA, USA). The samples were then incubated for 1 h at 37 °C, and heat-denatured at 95-98 °C for 5 min before being incubated on ice for 5 min. The samples were then divided into two tubes of 540 μL sample to which was added 540 μL of phenol/chloroform/isoamyl alcohol (25:24:1, pH=7.9) (ThermoFisher, MA, USA). The samples were gently mixed by inverting the tubes and centrifuged at 13,100 g for 10 min, before the aqueous (top) phase was transferred to a clean 2 mL tube. Glycoblue (ThermoFisher, MA, USA) was added to each aqueous phase (1:300 volume of aqueous phase), followed by 3 M, pH=5.2 sodium acetate (Carl Roth, DE) at 1:10 volume of the aqueous phase, after which 100% ethanol (VWR Chemicals, PA, USA) was added at 2.5x volume of the aqueous phase. Samples were then incubated at -20 °C for 1 h, before being centrifuged for 30 min, 13,100 g at 4 °C, and the supernatant removed. Pellets were washed in 500 μL ice-cold 70% ethanol (VWR Chemicals, PA, USA) and centrifuged for 10 min, 13,100 g at 4 °C. The pellets were dried in upside-down tubes till the ethanol had evaporated, at which point the still moist pellet was resuspended in 25 μL 10 mM Tris-HCl, pH=8 (ThermoFisher, MA, USA) and left to dissolve for 1 h at room temperature. The split sample fragments were then recombined into 50 μL of total DNA for experimental usage. DNA concentrations were measured using Qubit (ThermoFisher, MA, USA) as per manufacturer’s instructions.

### 2.4. Circular DNA extraction by SPRI bead purification method

Each 1 mL plasma sample was transferred into a 5 mL tube, and 10 μL of spike-in mix was added. Furthermore, a 1 mL negative H_2_O control was prepared for each purification. For each sample we added 50 μL of Proteinase K (>600 U/mL) (ThermoFisher, MA, USA), followed by 20 μl RNase A (7000 units/mL) (Qiagen, NL) and 750 μL of buffer 1 (containing an undisclosed (somewhere between 50-70% (w/w)) amount of Guanidinium Hydrochloride (EMD Millipore, DE) dissolved in ultra-pure H_2_O, supplemented with Tween 20 (Sigma-Aldrich, MA, USA), EDTA (Carl Roth, DE), and buffered by 10 mM Tris-HCl, pH=8 (ThermoFisher, MA, USA. The samples were then mixed by pipetting ≥10x till the solution was homogenous. Samples were next incubated for 30 min. at room temperature (15-25 °C). Afterwards, 1464 μL of AMPure XP beads (0.8x volume ratio) (Beckman Coulter, CA, USA) was added to each sample and mixed by pipetting 10x. 1710 μL buffer 2 (containing an undisclosed (between 75-90% (w/w)) guanidinium thiocyanate (Sigma-Aldrich, MA, USA) amount dissolved in isopropanol (Sigma-Aldrich, MA, USA)) was added to each sample which was mixed ≥10x until a homogenous solution was reached. The samples were then incubated at room temperature for 3 min. before the tubes were placed on a magnetic rack for 7 min. for bead-supernatant separation, after which the supernatant was extracted and discarded. The sample-beads were then washed with 3 mL 75-80% ethanol (VWR Chemicals, PA, USA) and again with 2 mL 75-80% ethanol. The supernatant was removed, and the samples were dried for a couple of minutes (ensuring that the beads were still lightly glossy). The tubes were then taken off the magnet, and the beads resuspended in 30 μL pH=8.0 salt-free elution buffer and mixed by pipetting 10x. The tubes were shortly spun down and incubated for 5 min. at 50 °C, before being placed on the magnet and incubated for two min. 25 μL of the eluate was taken out and transferred to a clean 1.5 mL DNA Lobind tube (Eppendorf, DE) (5 μL of the eluate was left behind). A second elution was then repeated using 25 μL elution buffer and combined with the former elute, leading to a final volume of 50 μL per sample.

To ensure an efficient qPCR, the purified product underwent an additional purification step in which 90 μL of AMPure XP beads (1.8x ration) (Beckman Coulter, CA, USA) was added to each sample, mixed by pipetting 10x and incubated at room temperature for 5 min. The samples were put on a magnetic rack for 3 min., following which the supernatant was discarded. The samples were then washed twice with 200 μL 75-80% ethanol (VWR Chemicals, PA, USA) and dried until the beads were slightly moist. The beads were then resuspended in 30 μL of elution buffer and mixed by pipetting 10x, before being incubated for 5 min. at 50 °C. This was followed by a 2 min. bead separation step on the magnetic rack, after which 25 μL of the eluate was taken out and transferred to a clean new 1.5 mL DNA Lobind tube (Eppendorf, DE) (5 μL were left behind). The elution step was then repeated using 27 μL elution buffer and combined with the other elute for a final sample volume of 52 μL. 12 μL of each sample were transferred to a new PCR tube for DNA concentration measurements using Qubit (ThermoFisher, MA, USA) as per manufacturer’s instructions, and qPCR quality control using SYBR™ Green PCR Master Mix (ThermoFisher, MA, USA) as per manufacturer’s instructions, targeting p4339 plasmid as an indicator of circular DNA preservation (**Table S1, Figure S1A,B**).

### 2.5. Enrichment of circular DNA by mitochondrial DNA linearization and linear DNA removal

Circular mtDNA was first linearized by adding 12.5 U/μL MssI enzyme (Pmel, 8-bp endonuclease) (ThermoFisher, MA, USA) for 2 h at 37 °C in accordance with the manufacturer’s instructions. Afterwards, the linear DNA was digested using 28.8 U/μL of plasmid-safe ATP-dependent DNase Exonuclease V (ExoV, RecBCD) (New England Biolabs, MA, USA) for 1 h at 37 °C, which was followed by a heat-inactivation step at 70 °C for 30 min as per manufacturer’s instructions. The circles of interest in the sample were then purified through the circle purification step detailed above (section 2.4.) using a 1.8x volume ratio of AMPure XP beads (Beckman Coulter, CA, USA). The efficiency of each sample purification was assessed through a standard qPCR assay using SYBR™ Green PCR Master Mix (ThermoFisher, MA, USA) as per manufacturer’s instructions, targeting Cytochrome c oxidase I (MT-CO1) as a control for mtDNA removal, the BCP1 yeast gene containing PCR fragment as an indicator of linear DNA digestion. Circular DNA preservation was assessed through qPCR targeting the p4339 plasmid (**Table S1, Figure S1B**).

### 2.6. Rolling-circle amplification of eccDNA for sequencing

The volume of each sample was reduced by 50% to ∼15 μL by evaporation at 55 °C for 1.5 h. The remaining sample was then used as a template for Φ29 polymerase reactions (4BB™ TruePrime® RCA Kit) (4basebio, UK), which was used in accordance with the manufacturer’s instructions and incubated for 48 h at 30 °C.

### 2.7. EccDNA sequencing from plasma samples

Laboratory A performed both the DNA fragmentation and library preparation for all samples. A portion from each Φ29-amplified DNA sample was diluted to 15 ng/μL in 10 mM Tris-HCl, pH=8 (ThermoFisher, MA, USA) for a total volume of 100 μL and sonicated (4 cycles of 20sec/30sec (on/off time)) using a Bioruptor (Pico II, Diagenode, BE). The successful generation of fragments with a mean fragment size of 400 nucleotides was confirmed using a Bioanalyzer as per manufacturer’s instructions (Agilent, CA, USA). The libraries were then prepared using NEBNext Ultra II DNA Library Prep Kit for Illumina and NEBNext Multiplex Oligos for Illumina (New England Biolabs, MA, USA) in accordance with the manufacturer’s protocol. The library replicate samples were prepared using the same procedure as applied to the original libraries, except only half the volume of reagents and amount of sonicated DNA was used.

Following library preparation, all samples were multiplexed and sequenced on a Novaseq 6000, S2 flow-cell as 2×150-nucleotide paired-end reads on two lanes (Rigshospitalet, DK), with an average of 124 million single reads per sample.

### 2.8. Mapping of the eccDNA with Circle-map

The sample sequence reads were mapped to a human reference genome (hg38) to record the origin of chromosomal-derived eccDNA (32). All steps were performed in accordance with GitHub instructions except for the usage of bwa mem, which was replaced by bwa mem2.

### 2.9. Circle quality assessment

Circles were deemed to be of sufficient quality and thereby trusted if they contained at least 1 concordant and 1 split read or 2 split reads while having at least 50% read coverage. Identified circles that did not fulfill these criteria were excluded from further analysis. Each uniquely mapped sequence fragment was counted as one circle. Circle sizes were calculated in bp using the end and start coordinates of the mapped circles.

### 2.10. Statistical analysis

The primary statistical and bioinformatics analysis was conducted using R-studio (V. 1.4.1106) and Ubuntu (20.04.2 LTS (GNU/Linux 4.4.0-19041-Microsoft x86_64)), which was applied for intersect analysis between circles and the ENSEMBL (Homo_sapiens.GRCh38.105.gtf.gz) genome. R-packages used in our analysis and graphical construction: rmarkdown, plyr, ggplot2, ggrepel, modelr, stringr, tidyr, tibble, tibble, sfsmisc, psych, car, quantreg, splines, tidyverse, tidyselect, writexl, readxl, rlang, dbplyr, dplyr, plotly, ggvenn, VennDiagram, BiocStyle::Biocpkg(“plotly”), RIdeogram (V.0.2.2), Bioconductor (V3.16), regioneR (V1.30.0), and pheatmap.

The genetic density and chromosome locations applied for ideogram formations were downloaded via the RIdeogram package from Gencode (version 32): gencode.v32.annotation.gff3.gz. The likelihood of achieving the observed number of eccDNA-overlaps for NFIA, PCDH9, ERBB4, and CTNND2 was assessed relative to randomized overlaps with the genetic regions from the same-sized theoretical eccDNA datasets (using regioneR (V1.30.0)) placed on a masked GR38 genome (BSgenome.Hsapiens.UCSC.hg38.masked (V1.4.5)).

Additional statistical testing: GraphPad Prism version 9.3.1 for Windows was applied for double-sided Student’s t-test comparisons. Figures were processed using Adobe Illustrator version 25.2.1.

## 3. Results

We have developed a SPRI bead purification method for eccDNA purification from human plasma. We compared our SPRI bead purification against a standard phenol/chloroform and salt precipitation method for the enrichment of eccDNA by using commercially available healthy plasma with spiked-in plasmids (**Figure S2**). The SPRI bead purification method led to a 35% greater DNA yield compared with the phenol/chloroform approach (p=0.0008, t-test), demonstrating that our method is more efficient than the phenol/chloroform and salt precipitation method.

Using SPRI bead purification, we tested the applicability of plasma eccDNA purification. One mL of plasma was analyzed per sample from four patients with stage IV lung cancer and four age-matched control donors, and eccDNA was purified in two independent laboratories (A and B) with replicates at four levels (**Figure 1A,B**). Following DNA extraction, the total DNA concentration, as well as the successful degradation of both linear DNA and mtDNA, were determined as methodological quality controls (**Figure S1**). One control plasmid (p4339) was also measured by qPCR to assess the method’s purification efficiency and the circular DNA loss during purification. For both laboratories A and B the plasmid was recovered 82.5% and 42.5%, respectively, and linear DNA and mtDNA were removed.

**Figure 1.**
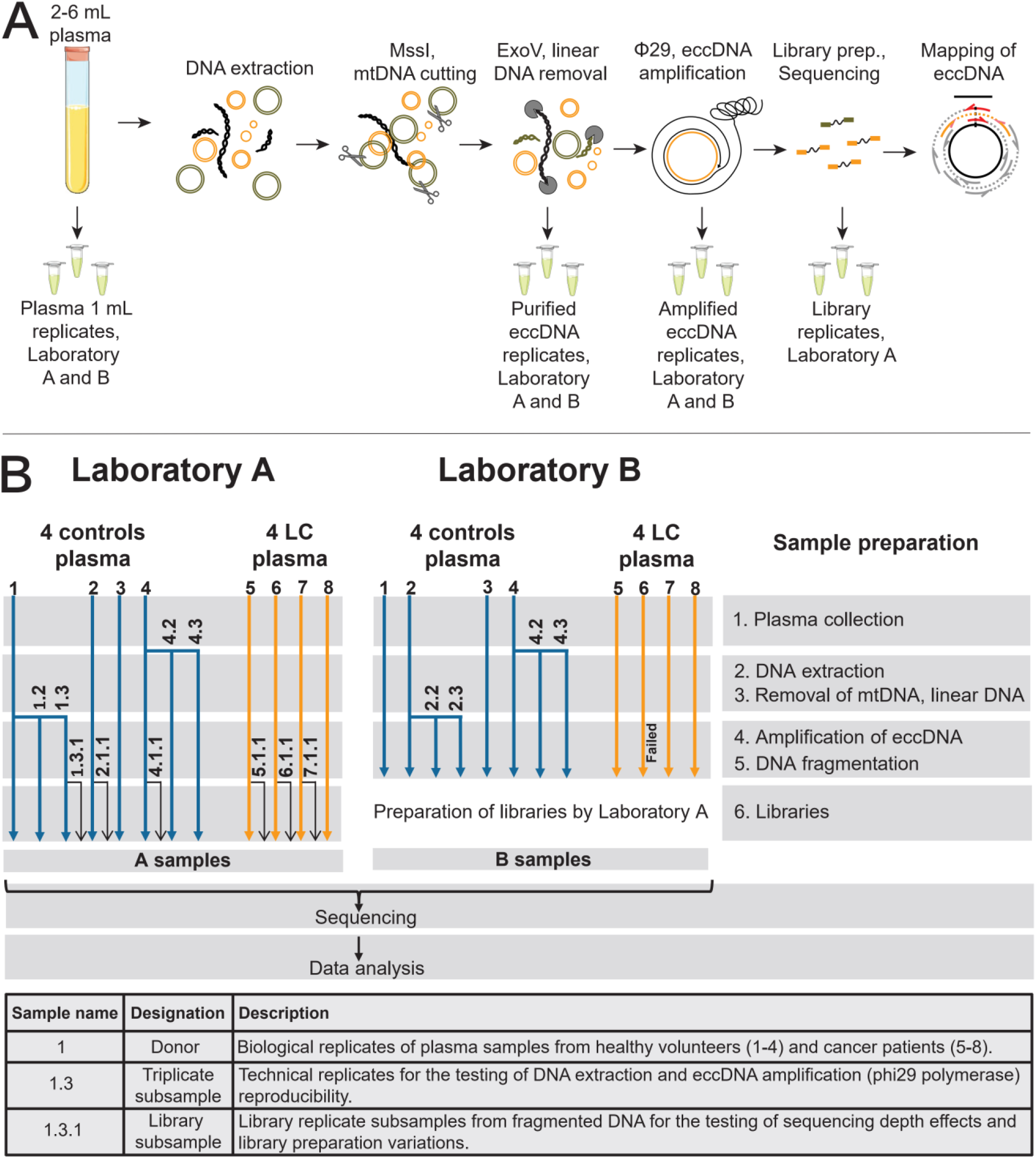
Design of the inter-laboratory comparison experiment. (A, B) At several steps during the workflow, technical replicates were subsampled to test the repeatability of the method. Samples 4, 4.2, 4.3 represent technical triplicates of a plasma sample, 1, 1.2, 1.3 and 2, 2.2, 2.3 represents Φ29 amplification replicates. Sample 6 in Laboratory B failed at Φ29 rolling-circle amplification step. Samples 1.3.1, 2.1.1, 4.1.1, 5.1.1, 6.1.1., and 7.1.1 were library preparation replicates only for Laboratory A samples, to assess divergence between samples of the new extraction method and sequencing. LC, lung cancer (adenocarcinoma); MssI, Pmel enzyme; ExoV, Exonuclease V; eccDNA, extrachromosomal circular DNA.

### 3.1. Experimental reproducibility

To evaluate the biological and technical variation of circular DNA, replicates were prepared in two different laboratories and at different stages of the analytical procedure following eccDNA purification (**Figure 1B**). The procedure was evaluated on plasma from different individuals within a biological group replicates from one individual, and technical replicates for the eccDNA purifications. As we are focusing on developing a new methodology, library preparation and sequencing was conducted in one laboratory only to avoid the introduction of any potential procedural variations. For each group and replicate, we compared the differences in eccDNA count and size (**Table 1**). We observed a high variance in eccDNA counts and sizes among all the samples from laboratories A and B, excluding library duplicates. SD for eccDNA counts were between 53% for controls (all the control A and B samples) and 43% for lung cancer samples (all the lung cancer A and B samples). SD for mean eccDNA size, excluding library duplicates, was 40% for controls (all the control A and B samples) and 29% for lung cancer samples (all the lung cancer A and B samples). This variation was as expected since the eccDNA counts varied markedly within the biological groups (SD for eccDNA counts were between 42% and 51% for controls (A1-A4 and B1-B4, respectively), and 35% and 52% for lung cancer (A5-A8 and B5-B8, respectively)). Also, a large SD was observed for the mean eccDNA size which was 26% and 27% for controls (A1-A4 and B1-B4, respectively) and 27% and 82% for lung cancer (A5-A8 and B5-B8, respectively).

**Table 1.** Variations of circle count and size among samples and between laboratories.

| Samples | Mean circle count | SD (%) | Mean circle size, bp | SD (%) | Median circle size, bp | Group | Variations |
| --- | --- | --- | --- | --- | --- | --- | --- |
| All the control A and B samples | 487 | 260 (53) | 3954 | 1570 (40) | 2212 | Controls | Between laboratories |
| All the lung cancer A and B samples | 2056 | 881 (43) | 2722 | 793 (29) | 1582 | Lung cancers | Between laboratories |
| A1 - A4 | 625 | 320 (51) | 3499 | 914 (26) | 1941 | Controls | Within individuals in a biological group |
| B1 - B4 | 402 | 168 (42) | 5277 | 1438 (27) | 3040 | Controls |  |
| A5 - A8 | 2278 | 787 (35) | 2530 | 678 (27) | 1562 | Lung cancers |  |
| B5 - B8 | 1813 | 949 (52) | 5610 | 4621 (82) | 4123 | Lung cancers |  |
| A4, A4.2, A4.3 | 583 | 200 (34) | 4098 | 538 (13) | 2261 | Controls | Within the same individual in each laboratory, triplicates |
| B4, B4.2, B4.3 | 537 | 121 (22) | 4949 | 1485 (30) | 2886 | Controls |  |
| A1, A1.2, A1.3 | 541 | 115 (21) | 1912 | 258 (14) | 1136 | Controls | Within the same eccDNA purification, triplicates |
| B2, B2.2, B2.3 | 246 | 28.3 (11) | 3652 | 196 (5) | 1769 | Controls |  |
| A1.3, A1.3.1 | 628 | 40 (6.3) | 1772 | 54 (3.1) | 1076 | Controls | Within the same library preparation |
| A2, A2.1.1 | 365 | 5 (1.2) | 4098 | 49 (1.2) | 2318 | Controls | Within the same library preparation |
| A4, A4.1.1 | 803 | 49 (6.1) | 3593 | 77 (2.1) | 1998 | Controls | Within the same library preparation |
| A5, A5.1.1 | 3117 | 256 (8.2) | 2195 | 111 (5) | 1385 | Controls | Within the same library preparation |
| A6, A6.1.1 | 2516 | 30 (1.2) | 3512 | 12 (0.3) | 2193 | Controls | Within the same library preparation |
| A7, A7.1.1 | 1636 | 122 (7.4) | 1806 | 68 (3.8) | 1286 | Controls | Within the same library preparation |

SD - standard deviation, bp – base pairs

Variation between individuals in a biological group was larger than between triplicate samples (**Table 1**). Triplicates of the same samples from the control group eccDNA count SD of 34% and 22% (A4, A4.2, A4.3 and B4, B4.2, B4.3, respectively), and eccDNA size SD of 13% and 30% (A4, A4.2, A4.3 and B4, B4.2, B4.3, respectively). The SD were the smallest for triplicates within the same purification, eccDNA count SD of 21% and 11%, and the eccDNA size SD of 14% and 5% (A1, A1.2, A1.3 and B2, B2.2, B2.3, respectively).

We also made duplicates during library preparations to test the variance within the same library preparation (**Table 1, Figure 2**). When comparing the eccDNA counts for samples sequenced from the same library preparation, we found little variation in eccDNA counts and size. For controls, SD of eccDNA count was 6.3% (A1.3 and A1.3.1), 1.2% (A2 and A2.1.1), and 6.1% (A4 and A4.1.1), for lung cancer SD was 8.2% (A5 and A5.1.1), 1.2% (A6 and A6.1.1), and 7.4% (A7 and A7.1.1). The size of eccDNA among the library triplicates again showed little variation when compared between the original samples and their corresponding library replicates. The SD for controls was 3.1% (A1.3 and A1.3.1), 1.2% (A2 and A2.1.1), and 2.1% (A4 and A4.1.1), for lung cancer samples, SD was 5% (A5 and A5.1.1), 0.3% (A6 and A6.1.1), and 3.8% (A7 and A7.1.1).

**Figure 2.**
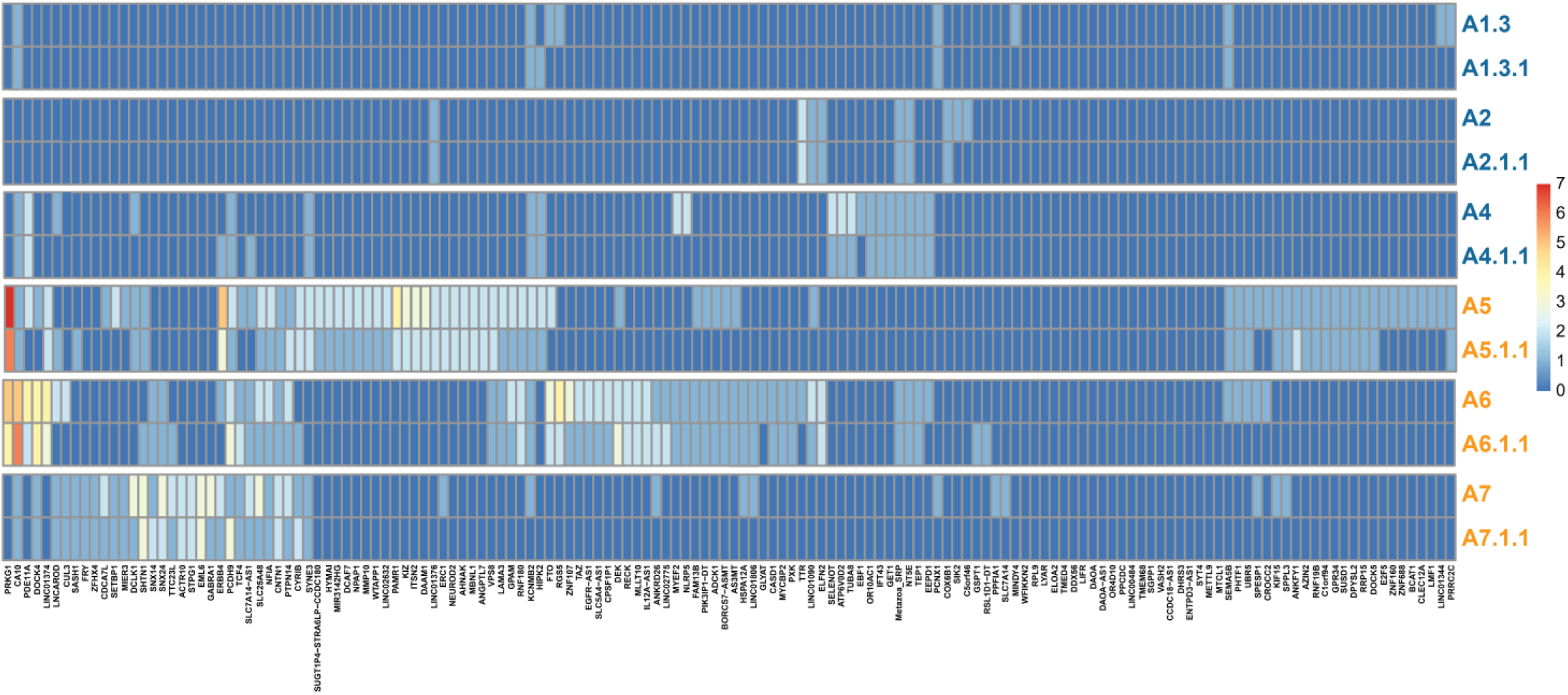
Heat map of the library duplicates. Reproducibility for 150 significantly overrepresented gene segments (>95 percentile quantile regression analysis) among all identified eccDNA relative to the genetic length. Controls n=3, cancers n=3. Blue letters, controls; orange letters, lung cancer samples.

Thus, though we found a large variation in the DNA sequence of eccDNA between individual samples, we were, to a large extent, able to reproduce the size distribution and the number of uniquely identified eccDNA among biological and technical replicates. On the other hand, we found large variations in the number and size of eccDNA detected in different laboratories, suggesting a need to formalize the protocol to reduce the variation.

### 3.2. Plasma from patients with lung cancer contains more eccDNA than healthy controls

Next, we analyzed the purified eccDNA in all samples obtained from both laboratories. We observed a significant difference in the mean unique eccDNA count between controls (487 circles) and lung cancer (2056 circles) samples for both laboratories (Laboratory A: p=0.0175, Laboratory B: p=0.0485) (**Figure 3A,B**). We did not find any significant difference in the mean eccDNA size between the two groups (range Laboratory A controls 67–109,902 bp and lung cancer 70–79,497 bp, range Laboratory B controls 33–64,091 bp and lung cancer 29–67,642 bp) (**Table S2**). As eccDNA may have diagnostic value, we then investigated the relative difference of eccDNA counts for each stage IV lung cancer sample relative to the control population (four age-matched control samples) from the same laboratory. Our Z-score analysis revealed that 3/4 of lung cancer samples from Laboratory A were significantly different (p<0.05) from the control population. When the same samples were tested at Laboratory B; all lung cancer samples (3/3) were significantly (p<0.01) different from the purified control samples (**Figure 3C**).

**Figure 3.**
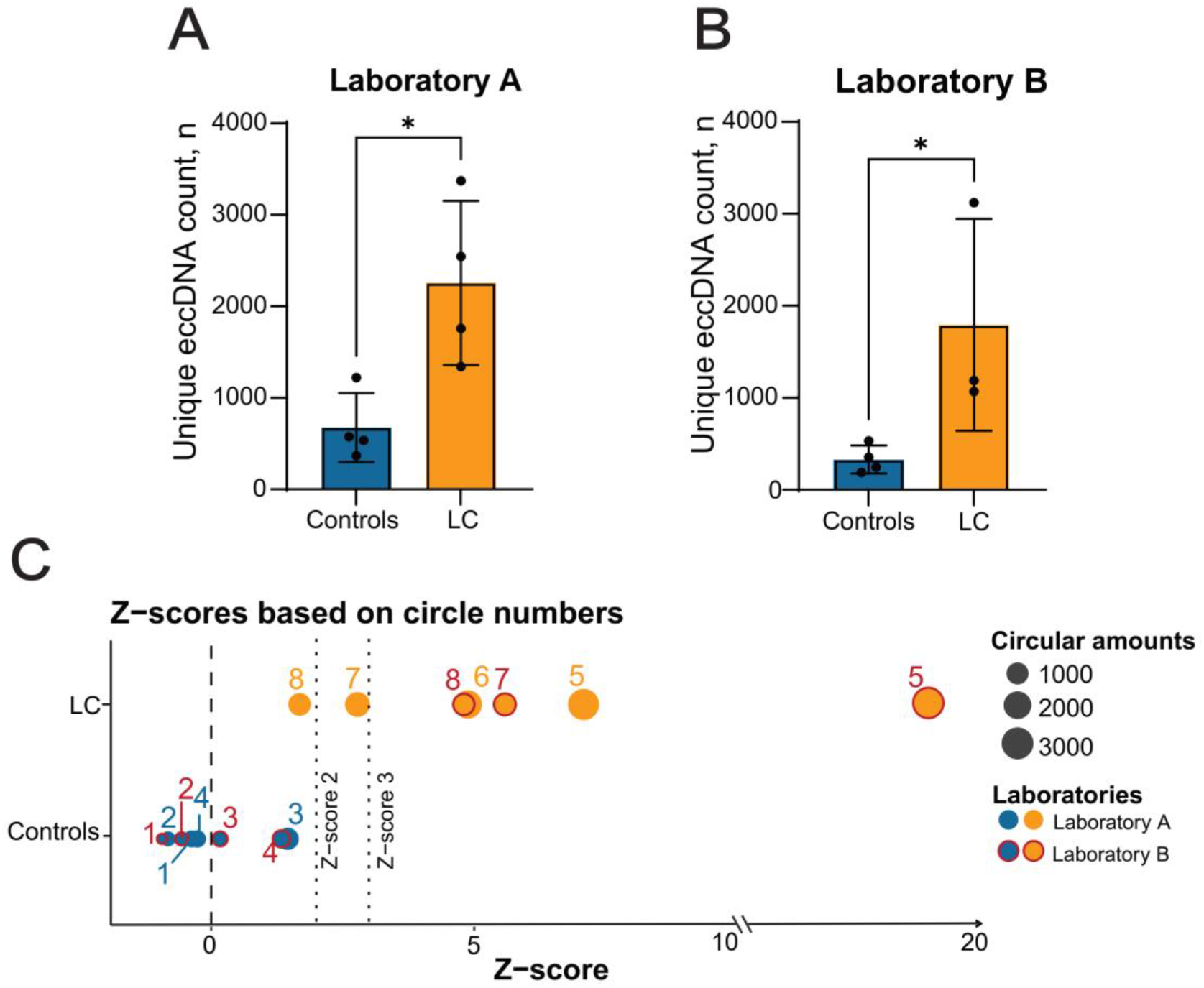
Circle count comparison between the control and lung cancer groups. (A, B) Number of unique eccDNA in the control and lung cancer groups in Laboratory A (controls n=4, cancers n=4) and B (controls n=4, cancers n=3). (C) Graphical presentation of the sample Z-scores relative to the mean control eccDNA counts of each laboratory. The mean of the sample triplicates were used for the Z-score assessments of the control values (counting as one assessment in the overall mean determination). Library duplicates are not included. LC, lung cancer (adenocarcinoma). *, P<0.05 α=0.05, SD. Dashed lines mark Z-scores of 0, 2 and 3.

### 3.3. EccDNA population characterization

An assessment of the investigated eccDNA size distribution revealed a periodic pattern in which the eccDNA sizes peaked at regular intervals of 170-200 bp (**Figure 4**). This pattern was also observed in plasma from both the control and the lung cancer group for circles containing gene segments (**Figure S3**).

**Figure 4.**
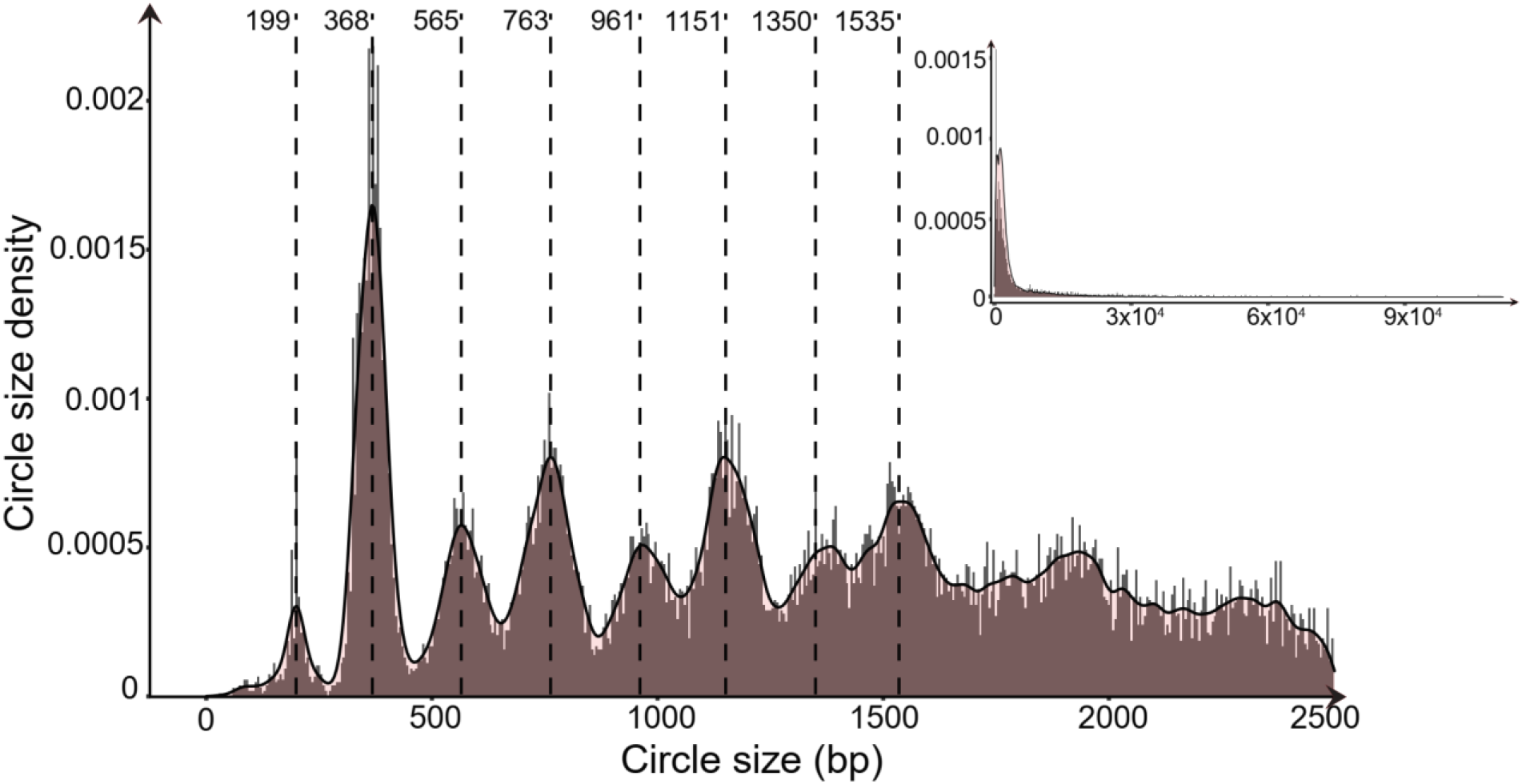
Combined eccDNA size density distributions for all eccDNA identified in this study. Dashed lines highlight peak density tips at 170-200 bp intervals observed in the plotted data. Controls n=8, cancers n=7. bp, base pairs.

EccDNA in somatic cells derives from gene-rich chromosomes (33). To test if this is also the case for eccDNA in plasma, we prepared ideograms of the eccDNA chromosomal distribution side-by-side with chromosomal gene density. We found that plasma eccDNA did not primarily originate from gene-rich chromosomes (such as chromosomes 17 and 19) but came from all parts of the genome (both healthy and lung cancer groups) (**Figure 5A,B**). We also assessed the relative eccDNA distribution per Mb against the genetic density per Mb for each chromosome (**Figure 5C**). In addition to having a higher eccDNA count, lung cancer samples were found to have a greater chromosomal variation (1.18 standard deviation (SD) (27.07%)) compared with control samples (0.31 SD (24.17%)).

**Figure 5.**
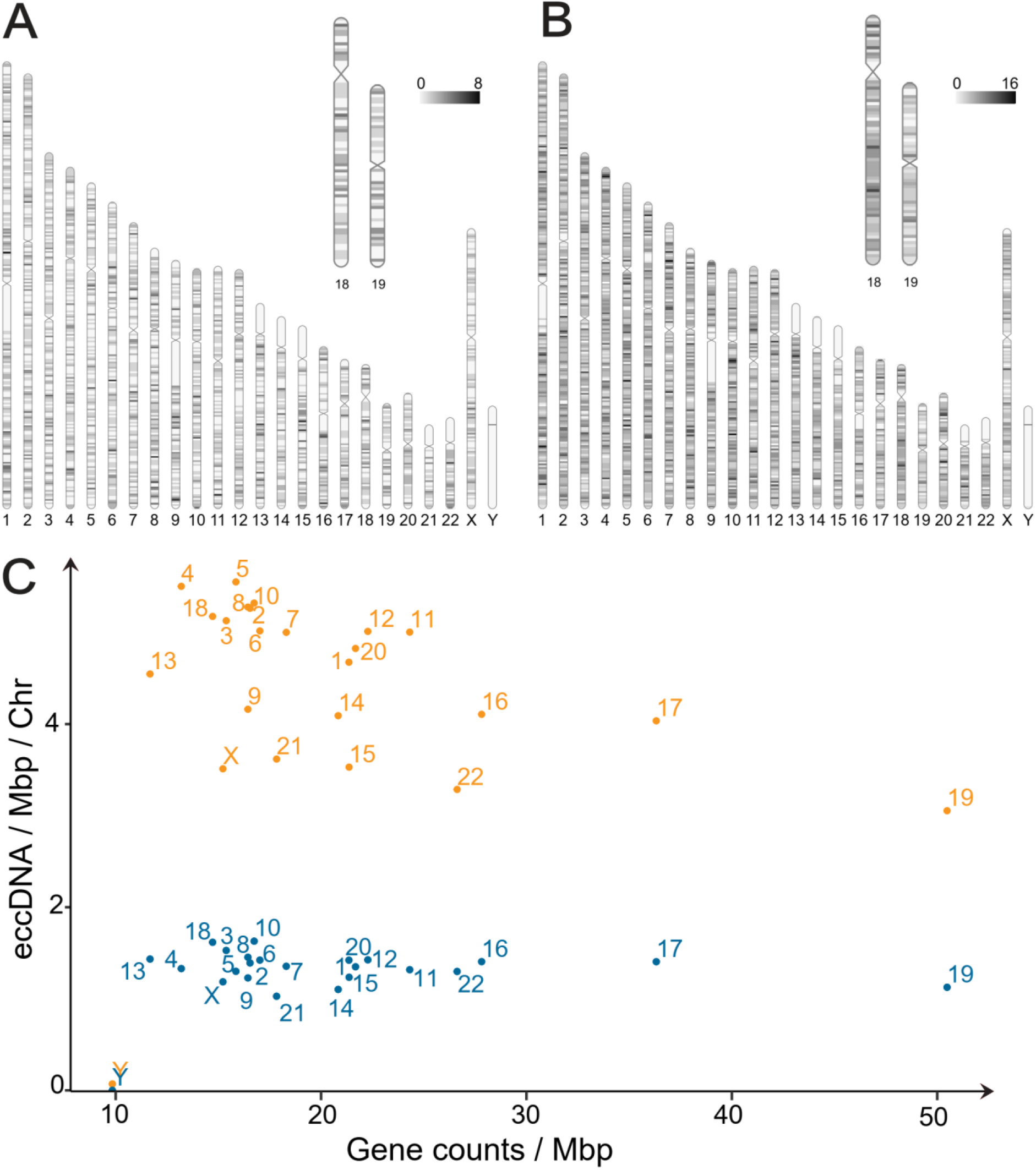
Chromosomal origin of eccDNA in controls and lung cancer patients. Ideograms of the chromosomal eccDNA distribution for mappable and non-mappable regions in one Mbp intervals (A) controls and (B) patients with lung cancer. Numeric range; controls: 0-8, lung cancer samples: 0-16. (C) Graphical overview of the chromosomal eccDNA/Mbp relative to gene counts/Mbp for all control and lung cancer samples. Controls n=8, cancers n=7. Chr, chromosome; Mpb, megabase pairs; blue dots, controls; orange dots, lung cancer samples.

### 3.4. The genetic origin of eccDNA

Genome coverage analysis showed that eccDNA purified from 1 mL plasma covered between 0.023-0.137% from healthy and 0.058-0.356% from lung cancer patients, as exemplified in a regional genome plot of eccDNA on chromosome 5 (**Figure 6**). A large portion of the plasma eccDNA did not contain any full genes or fragments of genes (9493 eccDNA in controls (40.25%) (n=19, including all the replicates and library duplicates) and 21268 in lung cancers (47.25%) (n=10, including all the replicates and library duplicates). The rest of the circles originated from genetic segments, and only a small fraction contained full-length genes (2.69% controls, 1.44% lung cancer), whereas most were found to contain fragments of genes (57.06% controls, 51.31% lung cancer). In general, we observed a low overlap in circles originating from the same genes among lung cancer samples and even less among controls (**Figure 7A,B; Table S3**). Despite the observed low overlap, the lung cancer genetic overlap was found to be greater than expected by chance (p=0.0099). Four genetic regions were particularly interesting, as fragments from these genes could be found in all cancer samples (NFIA, PCDH9, ERBB4, and CTNND2).

**Figure 6.**
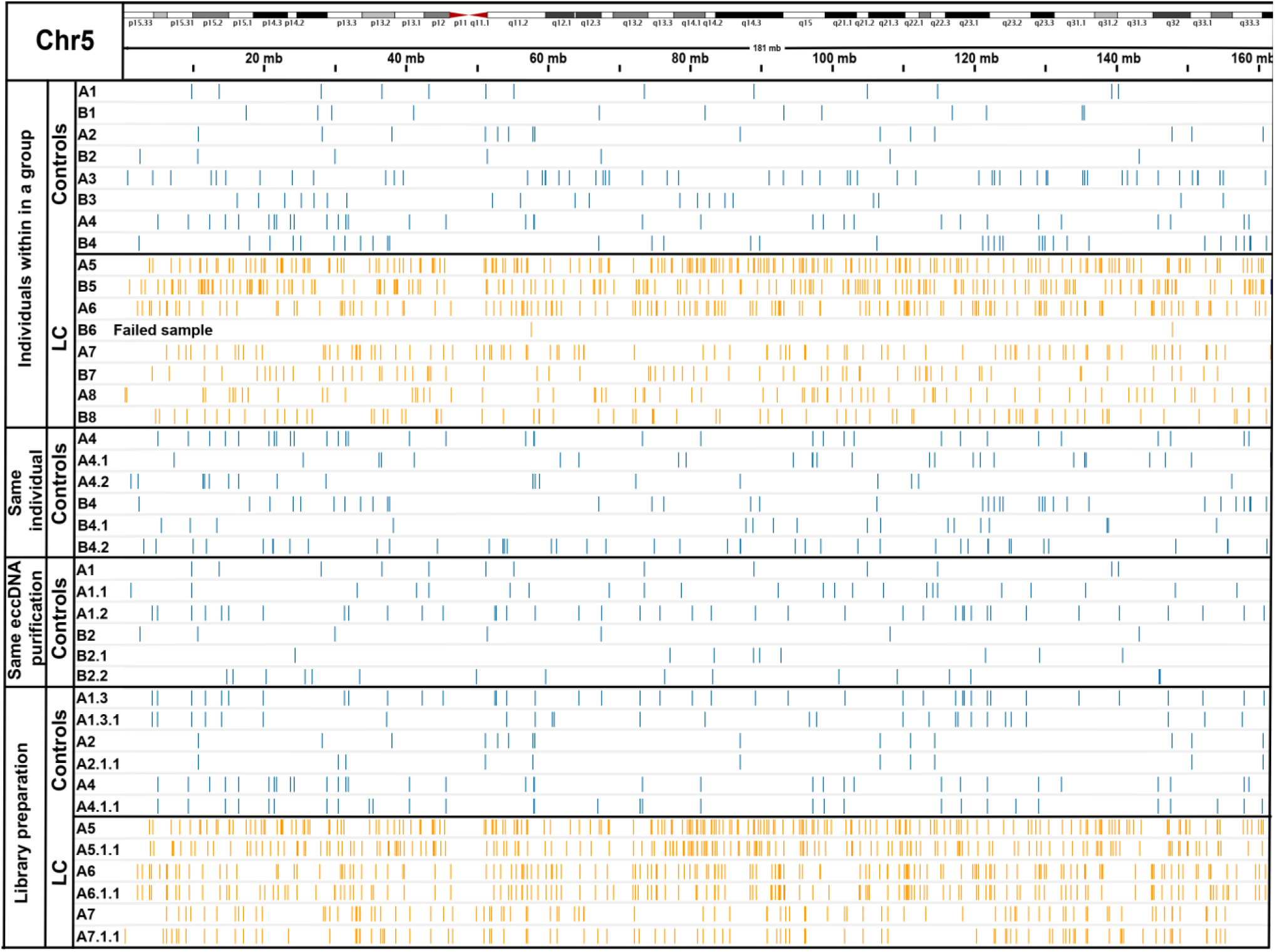
Coverage of eccDNA on Chr5 as an example. EccDNA covers a small fraction of the chromosome. Laboratory A (A1-A8) and B (B1-B8), each line represents a circle. Controls n=8, cancers n=7. LC, lung cancer (adenocarcinoma); blue lines, controls; orange lines, lung cancer samples.

**Figure 7.**
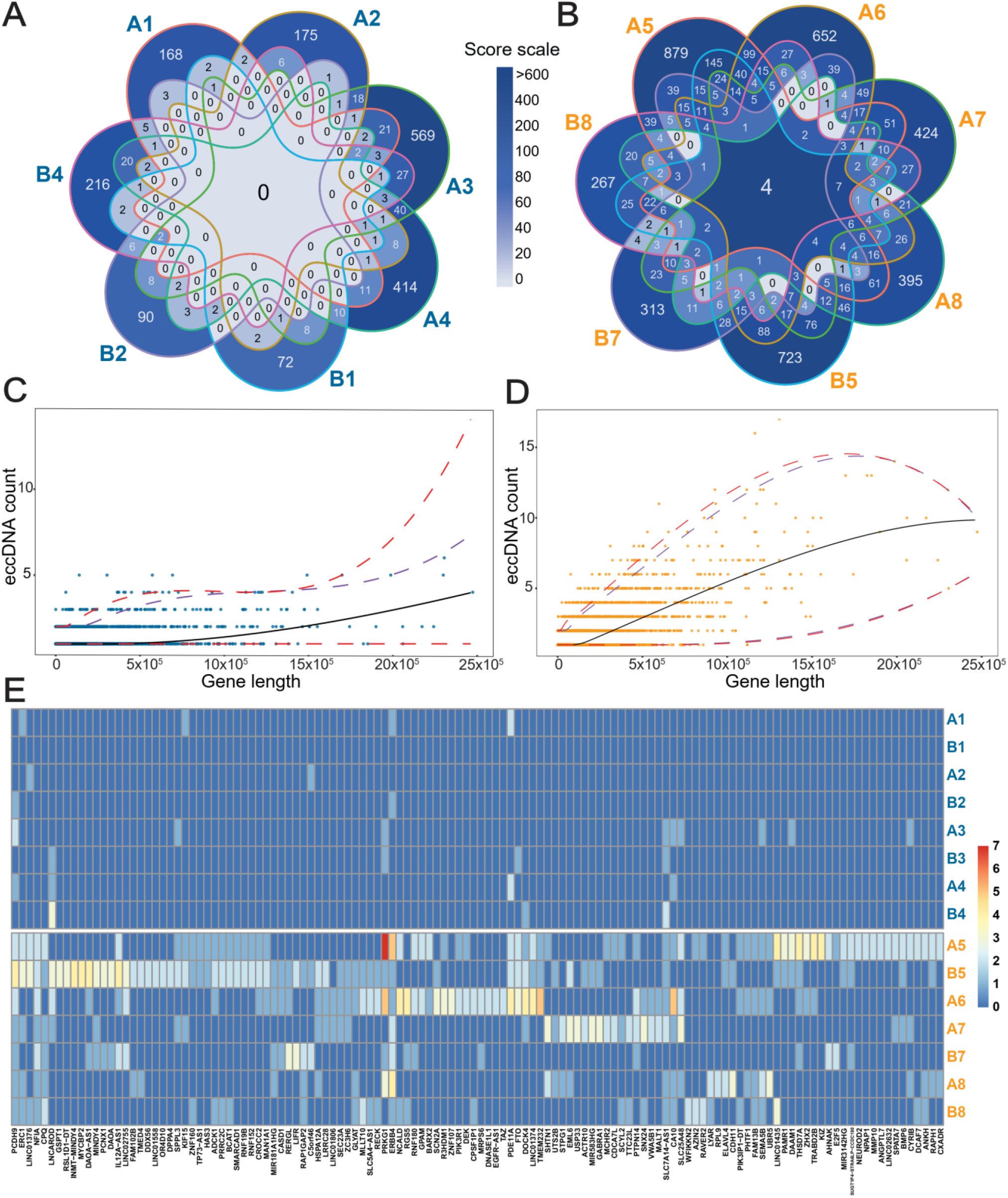
Reoccurring genes in eccDNA and the correlation between gene overlapping eccDNA count and gene length. (A) Gene overlapping eccDNA in controls (n=7) and (B) lung cancer samples (n=7). The formula for calculating the color intensity of frequency for overlapping gene fragments among the samples, Score scale=2 ^No. of samples in which a fragment was found^ x No. of common fragments of genes. For example, the center of lung cancer =27×4=512. (C, D) A quantitative regression analysis plot of a gene count found on eccDNA in correlation with the gene length.

We next assessed which of these genes were overrepresented on eccDNA relative to their lengths. To compensate for how larger genes may have a greater likelihood of generating eccDNA by chance, we conducted a quantitative regression analysis on gene length vs. eccDNA counts with 6 degrees of freedom and determined significant outliers (**Figure 7C,D**). Our analysis revealed 126 genes that were significantly overrepresented on eccDNA (>95% quantile) in the lung cancer samples. We then compared the sample profiles for these 126 genes on the eccDNA purified from control and lung cancer plasma samples and observed a low representation of these genes in the control samples (13.4x less hits pr. gene compared with lung cancer samples) (**Figure 7E; Table S4**). Gene ontology analysis of the 126 genes revealed an overrepresentation of genes involved in pathways important for cancer development, with the highest hits being in ontologies related to developmental growth and cell morphogenesis associated with differentiation (**Figure S4**).

Blue color, controls; orange color, cancer samples. Purple dashed lines mark 90% significance, red dashed lines mark 95% significance. (E) Heat map of the genes overrepresented on circles identified using quantitative regression analysis (C and D) in controls (top-blue) and cancer samples (bottom-orange) from laboratory A and B. Blue letters, controls; orange letters, lung cancer samples.

## 4. Discussion

In this study, we describe an effective method for the purification of eccDNA from plasma. We show that plasma from patients with stage IV lung cancer (adenocarcinoma) contains four times more unique eccDNA than healthy individuals, and that this number can be used to distinguish lung cancer patients from healthy controls (7/8 measures). As such, a large proportion of the unique eccDNA found in plasma from patients with stage IV lung cancer is, therefore, likely to be circulating tumor DNA (ctDNA). Through eccDNA sequence analysis, we observed that eccDNA originated from across the mappable parts of the genome. Notably, some gene regions were found to generate significantly more eccDNA compared with other regions. The eccDNA sequence analysis also reveals that eccDNA identified in one mL plasma covers less than 0.4% of the human genome, which can explain the large eccDNA sequence variations between samples from the same donor. Despite a significant variance in plasmid recovery between the two laboratories (Figure S1), the trends observed in circle count and genes found in the samples remained consistent.

We tested the biological and technical variation of eccDNA at various levels through comparison of the new SPRI bead purification method in two different laboratories. We observed a large variation in the DNA sequences present on the eccDNA among both samples from the same individual and eccDNA purifications from the same sample (Table S3). This variation is likely caused by the low coverage of the genome due to the low eccDNA content purifiable from the plasma. Our analysis showed that one mL of plasma only contained eccDNA, covering an average of 0.05% (control) and 0.17% (lung cancer) of the total genome (Figure 6). Thus, the replicative variations did not seem to originate from technical variations. The low amount of eccDNA however, can be problematic for the effective detection of eccDNA and might be overcome by larger plasma sample volumes. The observed variations in replication did not appear to stem from the tested methodology either, as it continued to diminish throughout the sampling process, and our library duplicates exhibited minimal variability. High biological variability for unique eccDNA among triplicates of different cell lines has been reported before using SPRI bead purification (34). As such, we consider the SPRI bead purification method to be reliable and reproducible.

Circulating eccDNA carries the potential as a ctDNA biomarker that can be used to complement studies of linear ctDNA. The amount of linear circulating cfDNA in the plasma of cancer patients and healthy individuals is low, and cell-free tumor DNA often represents only a small fraction of it (35,36). Though it is well established that linear cfDNA in plasma can be used to distinguish between healthy controls and individuals for several types of cancer, including different types of lung cancer (37–39), plasma eccDNA may provide an alternative and more stable biomarker for cancer detection. Cell-free eccDNA has only recently been identified and differentiated from linear cell-free DNA in the scientific community (rewieved in (26)). It could, therefore, potentially be used for early-stage diagnostics when ctDNA in plasma is very low, however, there is still a need for in-depth studies before it can be applied in a clinical setting.

Our results showed that the amount of unique eccDNA is significantly increased in plasma from lung cancer patients (Figure 3), which is similar to what has been observed for circulating linear cfDNA (35,36) and mentioned, though not further assessed, by Wu et al. for eccDNA in lung cancer patients (30). Interestingly, our findings suggest that this increase in eccDNA number can be applied as a potential marker for stage IV lung cancer. We observed no differences in the size of eccDNA between the lung cancer patients and the healthy controls. However, a previous study of plasma from four patients with lung cancer found two patients to carry larger circulating circular DNA pre-surgery than post-surgery (21).

We found that the new SPRI-bead purification method yields larger eccDNA compared to the methods used by Wu et al., Xu et al., and Kumar et al. Circular DNA in the current study stretches from 100 to >100.000 bp, whereas Wu et al. (100-400 bp) and Kumar et al. (100-1500 bp) show a far more limited detection range for their purified products (21,29,30). This suggests that the present method has a smaller loss of purified eccDNA than previously published methods despite using a far smaller sample volume.

We observed a high variation of the chromosomal plasma eccDNA load for lung cancer patients (Figure 5C), which may reflect an underlying chromosomal variation in the tumors that feed eccDNA into the plasma. Plasma eccDNA may therefore have the potential to reveal copy-number variations in the tumor, such as aneuploidy, which occurs in 90% of all solid cancers (reviewed in (40)).

In line with the low genome coverage observed for one mL plasma, we did not find any broad genetic-origin overlaps among eccDNA from controls, whereas in the lung cancer group, we found a number of eccDNAs from the same genes (Figure 7A,B). Some of the recurring genes were found to be involved in the regulation of developmental growth and differentiation, protein ubiquitination, and cancer-related pathways, as was also observed by Sanzhez-Vega et al. (41). Interestingly, fragments from four genes were found in all lung cancer samples and either at reduced levels, or not at all in the control populations. Plasma eccDNA containing NFIA, CTNND2, ERBB4, or PCDH9 are genes generally involved in cancer development (reviewed in (42–45)). An increased presence of these genes in the bloodstream could stem from genome amplifications and alterations within tumors. For instance, gene CTNND2 is located on Chr5, for which we demonstrated a greater eccDNA count per chromosome after length normalization in cancer samples (Figures 5C). Our assessments did not uncover any of the previously reported established biomarkers of lung cancer in our plasma eccDNA populations (5,29,30,46–50).

The 170-200 bp periodic peak size pattern observed among our purified plasma eccDNA (Figure 4) suggests that a large portion of the identified eccDNA has a nucleosomal-related origin. It has also been suggested that apoptotic or necrotic cells can lead to the release of nucleosomal-sized linear DNA (21,35). Likewise, similar periodical patterns have been observed for linear cfDNA in other cancers, peaking at 145 and 166 bp and with a high frequency of fragment sizes between 40 and 150 bp compared with cfDNA in healthy controls (51). The nucleosomal-related eccDNA pattern is also in line with the increased apoptosis of cancer cells, which leads to similar-sized fragments and can contribute to accelerated cancer development and metastasis (52). Furthermore, this periodical pattern was observed in both the lung cancer and control group (Figure S3), suggesting a general underlying eccDNA formation mechanism in which the nucleosomal structure plays a role for both cancers and healthy cells, as previously suggested for linear ctDNA (reviewed in (53)).

## 5. Conclusions

In conclusion, we observed a significant difference in plasma eccDNA counts between lung cancer and control samples. On the other hand, we were not able to identify conclusive gene markers that could be used to identify LC reproducibly. This suggests that traits other than genes and gene fragments on eccDNA are likely better targets for biomarker development (e.g. epigenetic features) due to the randomness of eccDNA found in plasma, its low genomic coverage, and its high inter-sample variability.

## Author’s contribution

EZ, LBH and BR designed the study. EZ and JH performed the purification of the eccDNA and sequence preparation. DG, MJA-B, mapped the circles based on sequence data. LBH and EZ performed bioinformatics and analysis. JSJ included the lung cancer patients in the LUCAS study and provided the samples and clinical data. EZ and LBH co-wrote the first draft of the manuscript. The article was prepared by EZ, BR and LBH. BR supervised the study. All authors discussed the results and contributed to the final manuscript.

## Ethics

All patients with Lung cancer gave written informed consent. The study was performed according to the declaration of Helsinki. The LUCAS study protocol was approved by the Ethics Committee of the Capital Region of Denmark (VEK j.nr. H-2-2011-1) and the Danish Data Protection Agency (j. nr. 2007-58-0015, HEH-750.24-56, I-suite nr. 02771 and PACTIUS P-2019-614).

## Acknowledgement

Professor Jørgen Wojtaszewski and technician Betina Blomgren, Department of Nutrition, Exercise and Sports, University of Copenhagen, for the sampling of blood and plasma from healthy individuals. Department of Oncology and The Danish Cancer Biobank for handling, freezing, and sampling the blood and plasma from patients with lung cancer. Sam Keating for processing test plasma samples with phenol/chloroform-salt precipitation method. Material from SMART-Servier Medical ART (https://smart.servier.com/) was used to create Figure 1.

