## Supplementary material for "Technical and biological variations in the purification of extrachromosomal circular DNA (eccDNA) and the finding of more eccDNA in the plasma of lung adenocarcinoma patients compared with healthy donors": Table S1

**Supplementary Table S1.** Description of primers for linear DNA fragment synthesis and qPCR reactions.

| **Gene** | **Primer sequence** | **Fragment length, bp** |
|---|---|---|
| GNP1 | F - GGTTCAAAGGTGTCGTTGCC | 337 |
| | R - GCACCGTTAGCAACGGAAAG | |
| AGP1 | F - GTTTTGGGTTTGCAGTCGCT | 820 |
| | R - GCACAGAAGGCAATAACGGC | |
| ACT1 | F - TGGATTCTGGTATGTTCTAGC | 1409 |
| | R - GAACGACGTGAGTAACACC | |
| BCP1 | F - TCAGTACAGTTGCGGTGGAC | 2716 |
| | R - TCGGATAGCCTCTGGTTAGG | |
| qPCR mtDNA | F - GCCCACTTCCACTATGTCCT | 92 |
|  | R GATTTTGGCGTAGGTTTGGTCT |  |
| qPCR BCP1 | F - CGGTGGTAACCCAGAAGTTGA | 130 |
| | R - TGTGGTGGTTGGGGAACCTA | |
| qPCR p4339 | F - TGCCCTGCCCCTAATCAGTA | 60 |
| | R - CTGGGCAGATGATGTCGAGG | |
