## Supplementary material for "Technical and biological variations in the purification of extrachromosomal circular DNA (eccDNA) and the finding of more eccDNA in the plasma of lung adenocarcinoma patients compared with healthy donors": Figure S2

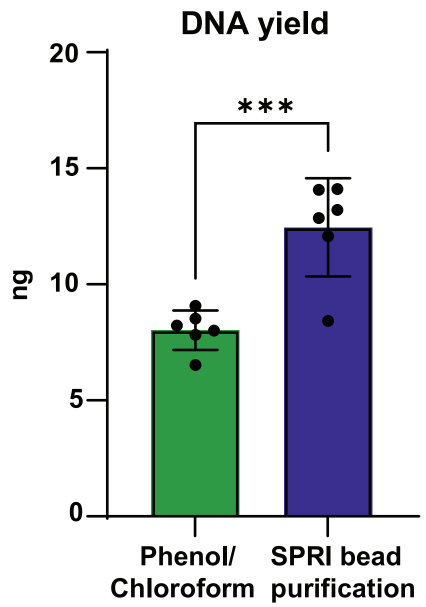


**Supplementary Figure S2.** Comparison of circular DNA yields from plasma from 6 technical replicates of healthy commercially available plasma. Circular DNA was purified with the Solid Phase Reversible Immobilization (SPRI) bead purification and phenol/chloroform-based salt precipitation method.
