## Supplementary material for "Technical and biological variations in the purification of extrachromosomal circular DNA (eccDNA) and the finding of more eccDNA in the plasma of lung adenocarcinoma patients compared with healthy donors": Table S2

**Supplementary Table S2.** eccDNA numbers and size differences among the samples and laboratories.
Sample characterization of eccDNA counts, sizes and Z-score relative to the mean control eccDNA counts of each laboratory. Library duplicates are not included.

### Laboratory A

| A samples | eccDNA numbers | Mean eccDNA Size, bp* | Median eccDNA Size, bp* | Z-score | Designation | Age, years | Days to Patient`s Death |
|---|---|---|---|---|---|---|---|
| **A1, A1.2, A1.3** | 536 | 1911.7 | 1041 | -0.37 | - | 60-65 | - |
| **A2** | 367 | 4048.9 | 2327 | -0.82 | - | 55-60 | - |
| **A3** | 1223 | 2371.7 | 1199 | 1.46 | - | 55-60 | - |
| **A4, A4.2, A4.3** | 576 | 4098.2 | 2365 | -0.26 | - | 50-55 | - |
| **A5** | 3373 | 2084.5 | 1317 | 7.17 | Lung cancer stage IV (adenocarcinoma) | 60 | 98 |
| **A6** | 2546 | 3500.2 | 2193 | 4.97 | Lung cancer stage IV (adenocarcinoma) | 63 | 102 |
| **A7** | 1757 | 1737.9 | 1221 | 2.88 | Lung cancer stage IV (adenocarcinoma) | 72 | 1795 |
| **A8** | 1343 | 2799.3 | 1518 | 1.77 | Lung cancer stage IV (adenocarcinoma) | 75 | 1492 |
| **A Laboratory Water** | 3 | 1893.3 | 2054 | -1.79 | - | - | - |

### Laboratory B

| B samples | eccDNA numbers | Mean eccDNA Size, bp** | Median eccDNA Size, bp** | Z-score | Designation | Age, years | Days to Patient`s Death |
|---|---|---|---|---|---|---|---|
| **B1** | 188 | 6288.5 | 3402.5 | -0.94 | - | 60-65 | - |
| **B2, B2.2, B2.1** | 244 | 3651.5 | 1743.5 | -0.57 | - | 55-60 | - |
| **B3** | 355 | 6754.3 | 4308 | 0.17 | - | 55-60 | - |
| **B4, B4.2, B4.3** | 531 | 4949.3 | 2184 | 1.33 | - | 50-55 | - |
| **B5** | 3120 | 3936.4 | 1725.5 | 18.47 | Lung cancer stage IV (adenocarcinoma) | 60 | 98 |
| **B6** | 12 | 13508.6 | 11668 | -2.1 | Lung cancer stage IV (adenocarcinoma) | 63 | 102 |
| **B7** | 1187 | 3147.6 | 1897 | 5.68 | Lung cancer stage IV (adenocarcinoma) | 72 | 1795 |
| **B8** | 1069 | 1847.4 | 1203 | 4.9 | Lung cancer stage IV (adenocarcinoma) | 75 | 1492 |
| **B Laboratory Water** | 4 | 2897.3 | 3179.5 | -2.15 | - | - | - |

*Laboratory A: t-test for controls vs lung cancer samples – mean size p=0.4337, median size p=0.6968
**Laboratory B: t-test for controls vs lung cancer samples – mean size p=0.9448, median size p=0.6553
