## Supplementary material for "Technical and biological variations in the purification of extrachromosomal circular DNA (eccDNA) and the finding of more eccDNA in the plasma of lung adenocarcinoma patients compared with healthy donors": Figure S3

A. Controls

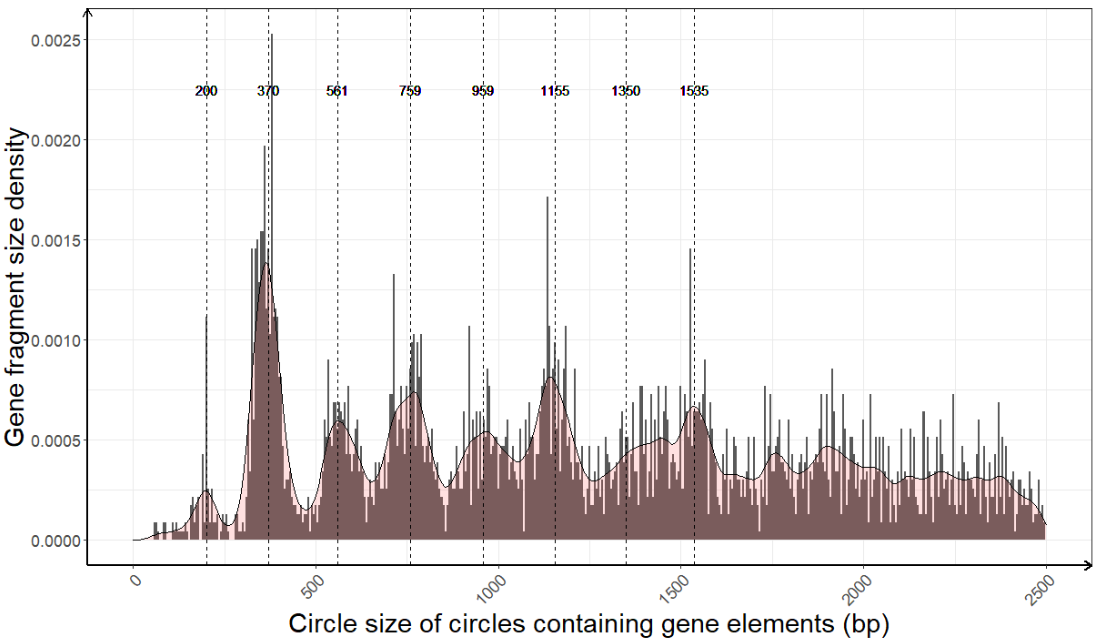


B. Lung cancer

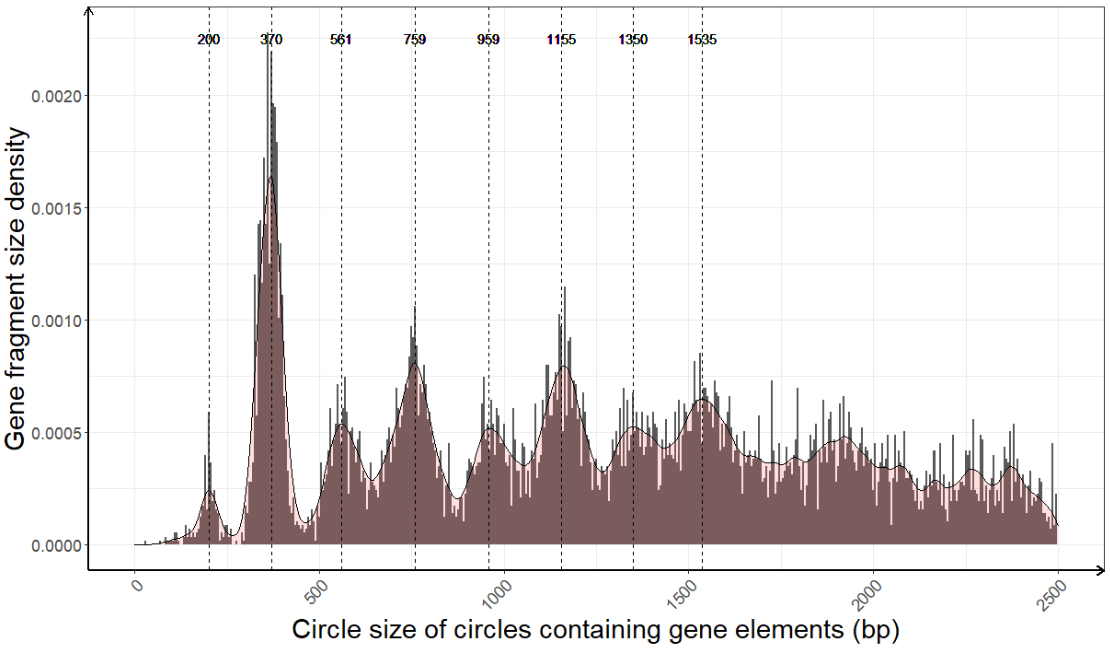


**Supplementary Figure S3.** eccDNA size density distributions of circles containing gene segments for (**A**) control samples (**B**) samples from patients with lung cancer (adenocarcinomas). Dashed lines highlight peak density tips at 170-200 bp intervals observed in the plotted data.
