## Supplementary material for "Technical and biological variations in the purification of extrachromosomal circular DNA (eccDNA) and the finding of more eccDNA in the plasma of lung adenocarcinoma patients compared with healthy donors": Table S3

**Supplementary Table S3**. Frequencies of co-occurring whole genes or gene fragments on eccDNA among samples.

| Groups | | Count of overlapping genes or gene fragments | | Percentage | |
|---|---|---|---|---|---|
| | | Average, range | | Average, range % | |
| | | Controls | Lung cancer | Controls | Lung cancer |
| Individuals within in a group | Lab A | 11.2<br>0-45 | 64.8<br>17-193 | 0.7<br>0-3% | 1.3<br>0-5% |
| | Lab B | 3.3<br>0-12 | 65.8<br>26-108 | 0.5<br>0-2% | 2.5<br>1-4% |
| Individuals between the two laboratories | Lab A/Lab B | 13.5<br>2-29 | 136<br>4-364 | 1.9<br>0.6-3.7% | 6.6<br>0.4-13.1% |
| Within the same individual | Lab A | 14.5<br>1-25 | - | 1.3<br>0-2% | - |
| | Lab B | 10.5<br>4-16 | - | 1<br>0-2% | - |
| Within the same eccDNA purification | Lab A | 10.5<br>2-17 | - | 1.3<br>0-2% | - |
| | Lab B | 3.3<br>0-9 | - | 0.8<br>0-2% | - |
| Within library preparation | Lab A | 252<br>170-352 | 929<br>626-1123 | 56.7<br>49.1-64.4% | 60.1<br>55.8-67.9% |
