## Supplementary material for "Technical and biological variations in the purification of extrachromosomal circular DNA (eccDNA) and the finding of more eccDNA in the plasma of lung adenocarcinoma patients compared with healthy donors": Figure S4

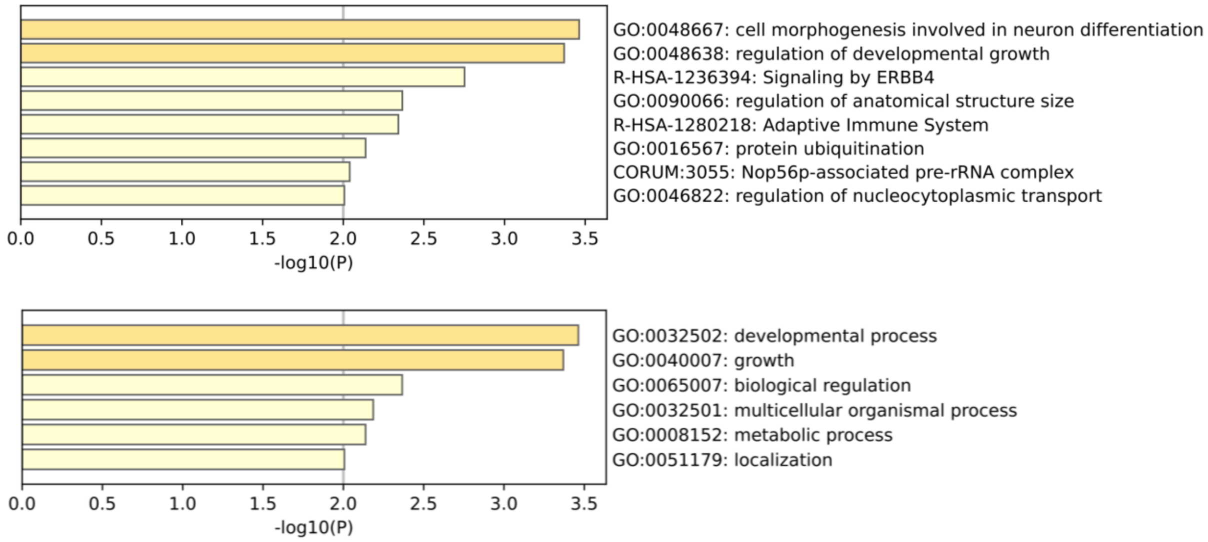


**Supplementary Figure S4.** Gene ontology analysis for the significantly overrepresented gene segments present on lung cancer eccDNA (Figure 7E, Supplementary Table S2). https://metascape.org.
