## Supplementary material for "Technical and biological variations in the purification of extrachromosomal circular DNA (eccDNA) and the finding of more eccDNA in the plasma of lung adenocarcinoma patients compared with healthy donors": Table S4

**Supplementary Table S4.** List of the recurring genetic fragments among eccDNAs found to be significantly overrepresented among cancer samples relative to the genetic length (identified through quantitative regression analysis, n=126, Figure 7E).

| Gene Name | Nr. of eccDNA | Gene Length, bp | Nr. of eccDNA /gene length | Quantile 0.025 | Quantile 0.05 | Quantile 0.5 | Quantile 0.95 | Quantile 0.975 | Relative 0.975 |
|---|---|---|---|---|---|---|---|---|---|
| TP73-AS1 | 2 | 11862 | 0.000169 | 0.999885 | 0.999977 | 0.99831 | 2.005558 | 1.991819 | 0.008181 |
| ANGPTL7 | 2 | 6626 | 0.000302 | 0.999912 | 0.999982 | 0.99864 | 1.997788 | 1.983032 | 0.016968 |
| RNF19B | 3 | 28364 | 0.000106 | 0.999942 | 0.999989 | 1.00148 | 1.996486 | 2.218852 | 0.781148 |
| AZIN2 | 3 | 42388 | 7.08E-05 | 1 | 1 | 1.00371 | 2.00939 | 2.497417 | 0.502583 |
| TRABD2B | 5 | 236857 | 2.11E-05 | 0.999738 | 0.999645 | 1.5519 | 4.028497 | 4.436887 | 0.563113 |
| ELAVL4 | 4 | 179743 | 2.23E-05 | 0.999885 | 1.000008 | 1.26272 | 3.520618 | 3.808382 | 0.191618 |
| NFIA | 11 | 597529 | 1.84E-05 | 0.98902 | 1.005049 | 3.42687 | 7.356415 | 8.226029 | 2.773971 |
| RAVER2 | 3 | 88137 | 3.40E-05 | 0.999507 | 0.999898 | 0.99691 | 2.476833 | 2.999315 | 0.000685 |
| PHTF1 | 3 | 62658 | 4.79E-05 | 0.999861 | 0.999969 | 0.99917 | 2.153909 | 2.787848 | 0.212152 |
| PRRC2C | 4 | 107981 | 3.70E-05 | 0.999338 | 0.999876 | 1.01508 | 2.749794 | 3.135055 | 0.864945 |
| PTPN14 | 6 | 203749 | 2.94E-05 | 1.000012 | 0.999952 | 1.38197 | 3.735408 | 4.071009 | 1.928991 |
| LINC02632 | 2 | 10441 | 0.000192 | 0.999889 | 0.999978 | 0.9983 | 2.004341 | 1.985456 | 0.014544 |
| PRKG1 | 17 | 1307535 | 1.30E-05 | 1.407526 | 1.486773 | 6.88765 | 12.9899 | 13.57549 | 3.424513 |
| LINC01374 | 7 | 371848 | 1.88E-05 | 0.994454 | 0.997202 | 2.25087 | 5.254932 | 5.903844 | 1.096156 |
| LINC01435 | 8 | 502876 | 1.59E-05 | 0.98889 | 0.998272 | 2.9339 | 6.473715 | 7.276075 | 0.723925 |
| GPAM | 4 | 65512 | 6.11E-05 | 0.999825 | 0.999961 | 0.99829 | 2.184482 | 2.817766 | 1.182234 |
| SHTN1 | 6 | 245109 | 2.45E-05 | 0.999583 | 0.999528 | 1.59438 | 4.102089 | 4.527774 | 1.472226 |
| LINC02755 | 9 | 653450 | 1.38E-05 | 0.992093 | 1.012619 | 3.71694 | 7.874157 | 8.767994 | 0.232006 |
| PAMR1 | 4 | 98477 | 4.06E-05 | 0.999393 | 0.99988 | 1.0032 | 2.621791 | 3.068017 | 0.931983 |
| AHNAK | 4 | 122693 | 3.26E-05 | 0.999348 | 0.999894 | 1.04527 | 2.933164 | 3.251393 | 0.748607 |
| MMP10 | 2 | 10126 | 0.000198 | 0.99989 | 0.999978 | 0.99831 | 2.003989 | 1.984427 | 0.015573 |
| BARX2 | 3 | 76431 | 3.93E-05 | 0.999671 | 0.99993 | 0.99585 | 2.317115 | 2.915032 | 0.084968 |
| ERC1 | 8 | 505424 | 1.58E-05 | 0.98883 | 0.998373 | 2.94719 | 6.497514 | 7.302141 | 0.697859 |
| BCAT1 | 4 | 139077 | 2.88E-05 | 0.999458 | 0.999933 | 1.09361 | 3.119061 | 3.396578 | 0.603422 |
| SCYL2 | 3 | 74575 | 4.02E-05 | 0.999698 | 0.999935 | 0.9961 | 2.293117 | 2.900116 | 0.099884 |
| SPPL3 | 4 | 141848 | 2.82E-05 | 0.999484 | 0.999941 | 1.10312 | 3.148847 | 3.422548 | 0.577452 |
| PCDH9 | 12 | 927611 | 1.29E-05 | 1.061394 | 1.107487 | 5.11026 | 10.29671 | 11.16975 | 0.830253 |
| DAAM1 | 4 | 182759 | 2.19E-05 | 0.999911 | 1.000007 | 1.27721 | 3.54812 | 3.840908 | 0.159092 |
| ADCK1 | 5 | 134905 | 3.71E-05 | 0.999423 | 0.999922 | 1.07998 | 3.073361 | 3.358208 | 1.641792 |
| SPATA7 | 3 | 85426 | 3.51E-05 | 0.999543 | 0.999905 | 0.99618 | 2.438955 | 2.980896 | 0.019104 |
| NPAP1 | 2 | 7618 | 0.000263 | 0.999905 | 0.999981 | 0.9985 | 2 | 1.981489 | 0.018511 |
| FTO | 7 | 456820 | 1.53E-05 | 0.990401 | 0.997075 | 2.69366 | 6.043821 | 6.80062 | 0.19938 |
| HAS3 | 2 | 13066 | 0.000153 | 0.999883 | 0.999976 | 0.99836 | 2.006149 | 1.999328 | 0.000672 |
| NEUROD2 | 2 | 6241 | 0.00032 | 0.999916 | 0.999983 | 0.99871 | 1.996824 | 1.984065 | 0.015935 |
| CA10 | 9 | 529704 | 1.70E-05 | 0.988408 | 0.999536 | 3.07378 | 6.724274 | 7.549207 | 1.450793 |
| DCAF7 | 3 | 43802 | 6.85E-05 | 1 | 1 | 1.00366 | 2.014477 | 2.522993 | 0.477007 |
| RNF152 | 3 | 86180 | 3.48E-05 | 0.999533 | 0.999903 | 0.99635 | 2.449457 | 2.986063 | 0.013937 |
| ZNF160 | 3 | 36828 | 8.15E-05 | 0.999988 | 0.999998 | 1.0033 | 1.997352 | 2.389249 | 0.610751 |
| LINC01376 | 6 | 361616 | 1.66E-05 | 0.994974 | 0.997353 | 2.19765 | 5.160579 | 5.794318 | 0.205682 |
| R3HDM1 | 4 | 193815 | 2.06E-05 | 0.999984 | 0.999989 | 1.33169 | 3.647448 | 3.961478 | 0.038522 |
| PDE11A | 8 | 449533 | 1.78E-05 | 0.990701 | 0.996982 | 2.65566 | 5.975892 | 6.724672 | 1.275328 |
| RAPH1 | 4 | 140990 | 2.84E-05 | 0.999476 | 0.999939 | 1.10014 | 3.139671 | 3.414466 | 0.585534 |
| ERBB4 | 16 | 1163124 | 1.38E-05 | 1.231792 | 1.299922 | 6.24036 | 12.09106 | 12.80907 | 3.19093 |
| CROCC2 | 4 | 86981 | 4.60E-05 | 0.999522 | 0.999901 | 0.99656 | 2.460644 | 2.991513 | 1.008487 |
| KIZ | 4 | 120639 | 3.32E-05 | 0.999341 | 0.99989 | 1.04026 | 2.908565 | 3.234296 | 0.765704 |
| CXADR | 3 | 80536 | 3.73E-05 | 0.999612 | 0.999918 | 0.99565 | 2.371759 | 2.946228 | 0.053772 |

| | | | | | | | | | |
|---|---|---|---|---|---|---|---|---|---|
| PIK3IP1-DT | 3 | 45522 | 6.59E-05 | 0.999997 | 0.999999 | 1.00352 | 2.021813 | 2.552889 | 0.447111 |
| KIF15 | 4 | 111655 | 3.58E-05 | 0.999331 | 0.999878 | 1.02133 | 2.797201 | 3.162732 | 0.837268 |
| SEMA5B | 5 | 119523 | 4.18E-05 | 0.999337 | 0.999888 | 1.03764 | 2.895069 | 3.225118 | 1.774882 |
| SLC7A14-AS1 | 7 | 412354 | 1.70E-05 | 0.992411 | 0.996846 | 2.46181 | 5.629986 | 6.334298 | 0.665702 |
| SMARCAD1 | 3 | 83681 | 3.59E-05 | 0.999567 | 0.999909 | 0.99589 | 2.414773 | 2.968775 | 0.031225 |
| ANKH | 4 | 166978 | 2.40E-05 | 0.999756 | 0.999999 | 1.20378 | 3.401528 | 3.673087 | 0.326913 |
| RNF180 | 5 | 207026 | 2.42E-05 | 1.000011 | 0.999934 | 1.39871 | 3.764306 | 4.107265 | 0.892735 |
| PIK3R1 | 3 | 86065 | 3.49E-05 | 0.999534 | 0.999903 | 0.99632 | 2.447854 | 2.985277 | 0.014723 |
| TMEM232 | 7 | 449723 | 1.56E-05 | 0.990693 | 0.996984 | 2.65665 | 5.977663 | 6.726655 | 0.273345 |
| FAM13B | 4 | 114001 | 3.51E-05 | 0.99933 | 0.999881 | 1.02578 | 2.826893 | 3.180899 | 0.819101 |
| MIR3142HG | 3 | 48832 | 6.14E-05 | 0.999984 | 0.999996 | 1.00303 | 2.039314 | 2.606829 | 0.393171 |
| SLC25A48 | 8 | 310568 | 2.58E-05 | 0.997408 | 0.998304 | 1.93261 | 4.692807 | 5.243514 | 2.756486 |
| BMP6 | 4 | 155629 | 2.57E-05 | 0.999629 | 0.999977 | 1.15539 | 3.290942 | 3.556897 | 0.443103 |
| DEK | 3 | 40688 | 7.37E-05 | 0.999999 | 1 | 1.00368 | 2.004424 | 2.465465 | 0.534535 |
| MCHR2 | 3 | 75728 | 3.96E-05 | 0.999681 | 0.999932 | 0.99594 | 2.307968 | 2.909448 | 0.090552 |
| MAN1A1 | 4 | 172556 | 2.32E-05 | 0.999815 | 1.000005 | 1.22902 | 3.454161 | 3.731688 | 0.268312 |
| THSD7A | 7 | 461833 | 1.52E-05 | 0.990202 | 0.997152 | 2.71981 | 6.090571 | 6.852756 | 0.147244 |
| CDCA7L | 3 | 45004 | 6.67E-05 | 0.999998 | 1 | 1.00357 | 2.019474 | 2.544023 | 0.455977 |
| DOCK4 | 8 | 480297 | 1.67E-05 | 0.989537 | 0.997545 | 2.81613 | 6.26287 | 7.043987 | 0.956013 |
| E2F5 | 3 | 40004 | 7.50E-05 | 0.999998 | 1 | 1.00364 | 2.002766 | 2.452279 | 0.547721 |
| CPQ | 8 | 504412 | 1.59E-05 | 0.988853 | 0.998332 | 2.94191 | 6.488062 | 7.291791 | 0.708209 |
| NCALD | 7 | 438365 | 1.60E-05 | 0.991187 | 0.996885 | 2.59741 | 5.871859 | 6.607909 | 0.392091 |
| UBR5 | 5 | 160486 | 3.12E-05 | 0.999684 | 0.999987 | 1.17557 | 3.338897 | 3.606078 | 1.393922 |
| ZHX2 | 4 | 192855 | 2.07E-05 | 0.999979 | 0.999992 | 1.32689 | 3.638898 | 3.950941 | 0.049059 |
| CYRIB | 4 | 177536 | 2.25E-05 | 0.999864 | 1.000008 | 1.25224 | 3.500357 | 3.784702 | 0.215298 |
| SUGT1P4-STRA6LP-CCDC180 | 4 | 138790 | 2.88E-05 | 0.999456 | 0.999933 | 1.09264 | 3.11595 | 3.39391 | 0.60609 |
| MIR181A1HG | 4 | 159610 | 2.51E-05 | 0.999674 | 0.999986 | 1.17187 | 3.330321 | 3.597145 | 0.402855 |
| FAM102B | 3 | 84811 | 3.54E-05 | 0.999551 | 0.999906 | 0.99606 | 2.430411 | 2.976651 | 0.023349 |
| MLLT10 | 5 | 218984 | 2.28E-05 | 0.999959 | 0.999845 | 1.46004 | 3.869858 | 4.239584 | 0.760416 |
| LNCAROD | 6 | 304633 | 1.97E-05 | 0.99766 | 0.998425 | 1.90186 | 4.638786 | 5.179032 | 0.820968 |
| HSPA12A | 5 | 179059 | 2.79E-05 | 0.999878 | 1.000008 | 1.25946 | 3.514352 | 3.801032 | 1.198968 |
| GLYAT | 4 | 91548 | 4.37E-05 | 0.999465 | 0.999891 | 0.99832 | 2.524783 | 3.022029 | 0.977971 |
| OR4D10 | 2 | 6046 | 0.000331 | 0.999917 | 0.999983 | 0.99874 | 1.996313 | 1.984682 | 0.015318 |
| RERGL | 5 | 239238 | 2.09E-05 | 0.999697 | 0.999612 | 1.56415 | 4.049709 | 4.463125 | 0.536875 |
| MYCBP2 | 6 | 284620 | 2.11E-05 | 0.998442 | 0.998829 | 1.79828 | 4.457266 | 4.960964 | 1.039036 |
| DAOA-AS1 | 4 | 46626 | 8.58E-05 | 0.999993 | 0.999998 | 1.00339 | 2.027165 | 2.571395 | 1.428605 |
| DAOA | 4 | 25167 | 0.000159 | 0.999922 | 0.999985 | 1.00065 | 1.999098 | 2.159075 | 1.840925 |
| SEC23A | 3 | 77727 | 3.86E-05 | 0.999652 | 0.999926 | 0.99574 | 2.334151 | 2.92513 | 0.07487 |
| ACTR10 | 3 | 35556 | 8.44E-05 | 0.999983 | 0.999997 | 1.00309 | 1.996124 | 2.363462 | 0.636538 |
| PCNX1 | 5 | 207977 | 2.40E-05 | 1.00001 | 0.999929 | 1.40358 | 3.772689 | 4.117791 | 0.882209 |
| LRRC28 | 4 | 139367 | 2.87E-05 | 0.999461 | 0.999934 | 1.09459 | 3.122199 | 3.399278 | 0.600722 |
| GSPT1 | 3 | 47954 | 6.26E-05 | 0.999988 | 0.999997 | 1.00319 | 2.034251 | 2.592968 | 0.407032 |
| RSL1D1-DT | 3 | 43962 | 6.82E-05 | 1 | 1 | 1.00365 | 2.015107 | 2.52583 | 0.47417 |
| CDH11 | 4 | 182359 | 2.19E-05 | 0.999907 | 1.000007 | 1.27527 | 3.544484 | 3.836584 | 0.163416 |
| RAP1GAP2 | 5 | 282036 | 1.77E-05 | 0.998534 | 0.99888 | 1.78492 | 4.433904 | 4.93274 | 0.06726 |
| EML6 | 6 | 248526 | 2.41E-05 | 0.99951 | 0.999476 | 1.61198 | 4.132625 | 4.565368 | 1.434632 |
| ZC3H6 | 3 | 64466 | 4.65E-05 | 0.999838 | 0.999964 | 0.99861 | 2.173046 | 2.807052 | 0.192948 |
| SCN2A | 4 | 197317 | 2.03E-05 | 0.999999 | 0.999979 | 1.34931 | 3.678549 | 4 | 4.44E-16 |
| SLC5A4-AS1 | 4 | 73267 | 5.46E-05 | 0.999717 | 0.999939 | 0.99633 | 2.276514 | 2.889256 | 1.110744 |
| DPPA4 | 2 | 11431 | 0.000175 | 0.999886 | 0.999977 | 0.9983 | 2.005251 | 1.989596 | 0.010404 |
| IL12A-AS1 | 7 | 323254 | 2.17E-05 | 0.996841 | 0.998048 | 1.99839 | 4.808547 | 5.381045 | 1.618955 |

| | | | | | | | | | |
|---|---|---|---|---|---|---|---|---|---|
| UTS2B | 3 | 63368 | 4.73E-05 | 0.999852 | 0.999967 | 0.99895 | 2.161327 | 2.795495 | 0.204505 |
| LIFR | 5 | 133686 | 3.74E-05 | 0.999413 | 0.999919 | 1.07616 | 3.059808 | 3.347169 | 1.652831 |
| MIR583HG | 4 | 165404 | 2.42E-05 | 0.999739 | 0.999996 | 1.19682 | 3.386486 | 3.656716 | 0.343284 |
| C5orf46 | 3 | 25812 | 0.000116 | 0.999926 | 0.999986 | 1.00081 | 1.998513 | 2.170753 | 0.829247 |
| GABRA1 | 4 | 52918 | 7.56E-05 | 0.999959 | 0.99999 | 1.00211 | 2.066649 | 2.667317 | 1.332683 |
| LINC01558 | 2 | 12335 | 0.000162 | 0.999884 | 0.999977 | 0.99832 | 2.005836 | 1.994544 | 0.005456 |
| INMT-MINDY4 | 4 | 139944 | 2.86E-05 | 0.999466 | 0.999936 | 1.09655 | 3.128428 | 3.404663 | 0.595337 |
| MINDY4 | 4 | 120970 | 3.31E-05 | 0.999342 | 0.999891 | 1.04105 | 2.91255 | 3.237033 | 0.762967 |
| DDX56 | 2 | 9247 | 0.000216 | 0.999895 | 0.999979 | 0.99834 | 2.002838 | 1.982319 | 0.017681 |
| TMED4 | 3 | 4393 | 0.000683 | 0.999935 | 0.999987 | 0.99912 | 1.99133 | 1.992529 | 1.007471 |
| CASD1 | 3 | 47800 | 6.28E-05 | 0.999989 | 0.999997 | 1.00321 | 2.033394 | 2.590504 | 0.409496 |
| RECK | 4 | 87542 | 4.57E-05 | 0.999515 | 0.9999 | 0.99672 | 2.468495 | 2.995308 | 1.004692 |
| VWA5B1 | 3 | 68643 | 4.37E-05 | 0.999783 | 0.999953 | 0.99739 | 2.220192 | 2.848223 | 0.151777 |
| RGS5 | 6 | 77389 | 7.75E-05 | 0.999657 | 0.999927 | 0.99576 | 2.329687 | 2.92252 | 3.07748 |
| MALT1 | 3 | 83012 | 3.61E-05 | 0.999576 | 0.999911 | 0.99581 | 2.405556 | 2.964061 | 0.035939 |
| LINC01806 | 2 | 4320 | 0.000463 | 0.999936 | 0.999987 | 0.99914 | 1.991082 | 1.992986 | 0.007014 |
| MRPS6 | 3 | 69810 | 4.30E-05 | 0.999766 | 0.999949 | 0.99709 | 2.234029 | 2.858996 | 0.141004 |
| CPSF1P1 | 3 | 3729 | 0.000805 | 0.999943 | 0.999988 | 0.99931 | 1.988985 | 1.997036 | 1.002964 |
| SNX24 | 5 | 185915 | 2.69E-05 | 0.999935 | 1.000004 | 1.29255 | 3.57669 | 3.87513 | 1.12487 |
| EGFR-AS1 | 2 | 9184 | 0.000218 | 0.999895 | 0.999979 | 0.99835 | 2.002745 | 1.982211 | 0.017789 |
| ZNF107 | 3 | 45483 | 6.60E-05 | 0.999997 | 0.999999 | 1.00353 | 2.021633 | 2.552225 | 0.447775 |
| TAZ | 2 | 10202 | 0.000196 | 0.99989 | 0.999978 | 0.9983 | 2.004077 | 1.984662 | 0.015338 |
| DNASE1L1 | 2 | 10876 | 0.000184 | 0.999888 | 0.999977 | 0.99829 | 2.004777 | 1.987107 | 0.012893 |
| STPG1 | 3 | 59935 | 5.01E-05 | 0.999892 | 0.999976 | 1.00004 | 2.126695 | 2.757184 | 0.242816 |
| USP33 | 3 | 63865 | 4.70E-05 | 0.999846 | 0.999966 | 0.99879 | 2.166594 | 2.800766 | 0.199234 |
| TTC23L | 3 | 60623 | 4.95E-05 | 0.999885 | 0.999974 | 0.99982 | 2.13338 | 2.765137 | 0.234863 |
| LYAR | 3 | 22453 | 0.000134 | 0.999907 | 0.999982 | 0.99996 | 2.001666 | 2.112596 | 0.887404 |
| RPL9 | 2 | 6344 | 0.000315 | 0.999915 | 0.999983 | 0.99869 | 1.997088 | 1.983764 | 0.016236 |
| WFIKKN2 | 2 | 7703 | 0.00026 | 0.999904 | 0.999981 | 0.99848 | 2.000172 | 1.98143 | 0.01857 |

bp, base pairs
