## Supplementary material for "Technical and biological variations in the purification of extrachromosomal circular DNA (eccDNA) and the finding of more eccDNA in the plasma of lung adenocarcinoma patients compared with healthy donors": Figure S1

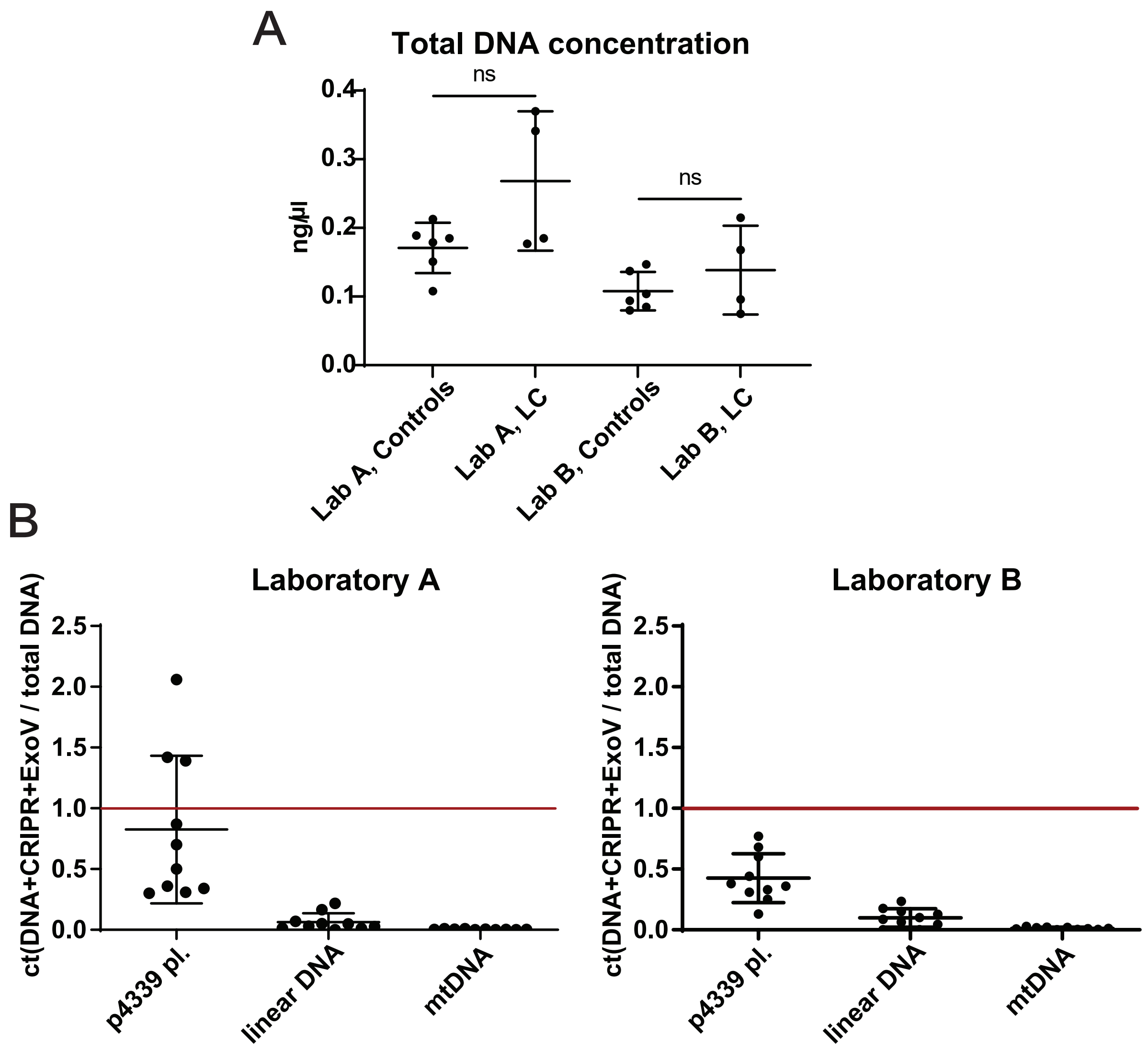


**Supplementary Figure S1.** (**A**) Total DNA concentration measured by Qubit after DNA extraction from plasma samples. (**B**) Quality tests for the extracted circular DNA measured by qPCR. The red bar indicates the total amount of DNA before MssI and Exonuclease V (ExoV) treatment. Ct, cycle threshold; LC, lung cancer, p4339, iner control plasmid; ns, not significant.
